# Spatiotemporal transcriptomics reveals pathogenesis of viral myocarditis

**DOI:** 10.1101/2021.12.07.471659

**Authors:** Madhav Mantri, Meleana M. Hinchman, David W. McKellar, Michael F. Z. Wang, Shaun T. Cross, John S. L. Parker, Iwijn De Vlaminck

**Affiliations:** Nancy E. and Peter C. Meinig School of Biomedical Engineering, Cornell University, Ithaca, New York; Baker Institute for Animal Health, College of Veterinary Medicine, Cornell University, Ithaca, New York; Cornell Institute for Host-Microbe Interactions and Disease, Cornell University, Ithaca, New York

## Abstract

A significant fraction of sudden death in children and young adults is due to myocarditis, an inflammatory disease of the heart, most often caused by viral infection. Here we used integrated single-cell and spatial transcriptomics to create a high-resolution, spatially resolved map of reovirus-induced myocarditis in neonatal murine hearts. We assayed hearts collected at three timepoints after reovirus infection and studied the temporal, spatial, and cellular heterogeneity of host-virus interactions. We further assayed the intestine, the primary site of reovirus infection to establish a full chronology of molecular events that ultimately lead to myocarditis. We implemented targeted enrichment of viral transcripts to establish the cellular targets of the virus in the intestine and the heart. Our data give insight into the cell-type specificity of innate immune responses, and into the transcriptional states of inflamed cardiac cells in reovirus-infected heart. We find that inflamed endothelial cells recruit cytotoxic T cells and undergo pyroptosis in the myocarditic tissue. Analyses of spatially restricted gene expression in myocarditic regions and the border zone around those regions identified immune-mediated cell-type specific injury and stress responses. Overall, we observe a dynamic and complex network of cellular phenotypes and cell-cell interactions associated with viral myocarditis.

## INTRODUCTION

Viral infection is the most common cause of myocarditis^1,2^. The resulting inflammatory cardiomyopathy can lead to arrhythmias, dilated cardiomyopathy, and death^1,3,4^. In humans, viral myocarditis is challenging to study because of the low sensitivity of available diagnostic testing, the acute onset of the disease, the focal nature of the disease, and the extreme heterogeneity of immune-virus interactions in complex cardiac tissues^4–6^. In mice, mammalian orthoreovirus offers a flexible model system^7^. After oral inoculation, the Type 1 Lang (T1L) reovirus strain initially infects the gastrointestinal tract. Within days the infection then spreads to secondary sites in the body, including the heart, leading to myocarditis in up to 50% of infections^7–9^. Yet, even in this mouse model, the molecular pathogenesis of viral myocarditis is difficult to study because of the complex network of cardiac and immune cell types involved and the cellular, spatial, and temporal heterogeneity of the disease^2,10^. Consequently, neither the cell types that are responsible for the innate immune response, nor the cell types that are infected *in vivo* have been identified. Similarly, the responses of infected and uninfected bystander cells within the heart have not been characterized. In addition, the protective versus damaging effects of adaptive immune responses have not been quantified. Experiments in mice with severe combined immunodeficiency (SCID) indicate that adaptive immune responses are not required for myocardial injury and heart failure^7,11^, but these observations do not exclude the possibility that immune-cell-mediated injury is important in immunocompetent mice. Unbiased characterization of all cellular phenotypes as a function of time and location within infected cardiac tissues is needed to address these knowledge gaps.

Here we used integrated single-cell and spatially resolved RNA-sequencing (RNA-seq) to study the cellular and spatial heterogeneity of myocarditic processes in the hearts of reovirus-infected neonatal mice at multiple time points after infection. We also applied these technologies to study the innate response to reovirus infection in the intestine. In addition, we performed time-series single-cell RNA-seq (scRNA-seq) of cardiac tissues of mice infected with a reovirus point mutant that does not cause myocarditis. To establish viral tropism, we implemented molecular enrichment of non-polyadenylated viral transcripts that were otherwise poorly represented in the transcriptomes. Our measurements give insight into the cell-type specificity of innate immune responses, into the tropism of the virus in the intestine and the heart, and into the transcriptional states of cell types involved in the production of inflammatory cytokines and the recruitment of circulating immune cells. Analyses of spatially restricted gene expression in myocarditic regions and the border zone around those regions identified injury and stress responses in different cell types, including cardiomyocytes. Overall, our data identify spatially restricted cellular interactions and cell-type specific host responses during reovirus-induced myocarditis.

## RESULTS

### Single-cell and spatial transcriptomics of reovirus T1L-infected neonatal mice hearts

To elucidate the pathogenesis of reovirus-induced myocarditis, we analyzed heart tissues collected from neonatal mice infected orally with either the T1L strain of reovirus or a mock control (**Methods, Fig. 1A**). We generated scRNA-seq data for 31,684 cells from infected hearts and mock controls at 4, 7, and 10 days post-infection (dpi), and 8,243 spatial transcriptomes for four tissue sections from infected hearts and mock controls at 4 and 7 dpi from the same litter (10x Chromium and 10x Visium, **Methods, Supp Fig. 1A-1B and Fig. 1B-1C**). The single-cell transcriptomes represented 18 distinct cell types, including cardiomyocytes, endocardial cells, cardiac fibroblasts, endothelial cells, mural cells, macrophages, neutrophils, NK cells, dendritic cells, T cells, and B cells (**Methods, Fig 1B, Supp Data 1, Supp Fig. 1C-F**). Clustering of the spatial transcriptomic data revealed distinct transcriptional programs for myocarditic regions and the border zone surrounding the myocarditic regions in the 7 dpi reovirus-infected heart that corresponded to areas of tissue damage identified by H&E staining (**Fig. 1C, Supp Fig. 2A-B**). The combination of scRNA-seq and spatial transcriptomics allowed us to resolve and visualize cell types and gene expression in a spatial context (**Supp Fig. 2C**). Because the virus first infects the gastrointestinal tract before it spreads to other body sites including the heart, we also performed scRNA-seq and spatial transcriptomics on ileum. We obtained 7,695 single-cell transcriptomes and 8,027 spatial spot transcriptomes for ileum from mock and infected samples at 1 and 4 dpi (**Fig. 1D, Supp Fig. 3A-D**).

**Figure 1:**
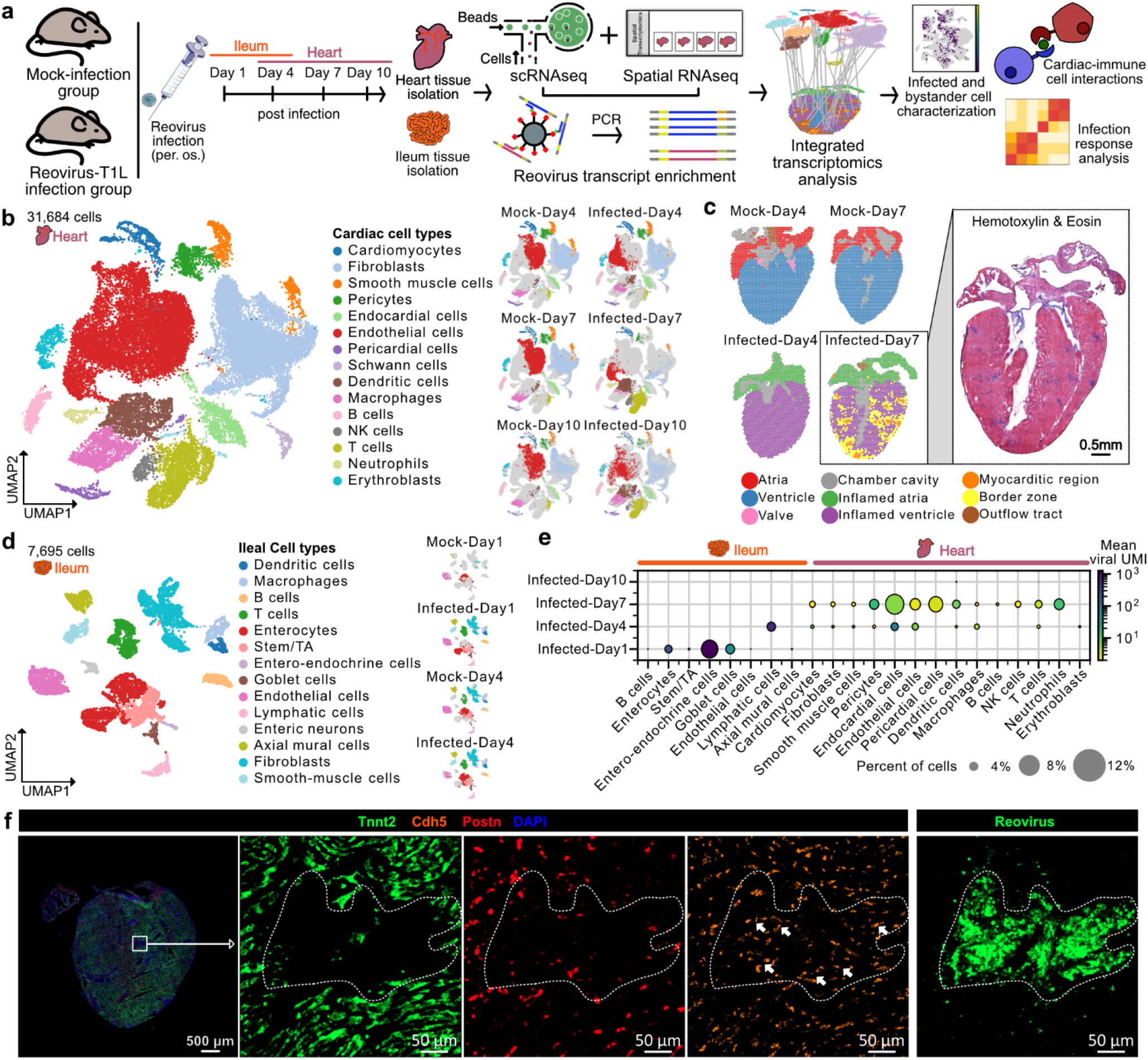
Single-cell and spatial transcriptomics of cardiac and ileum tissue of reovirus-infected neonatal mice. **a)** Experiment and analysis workflow. **b)** UMAP plot of 31,684 single-cell transcriptomes from mock-infected and reovirus-infected hearts at 4, 7, and 10 dpi (one animal per condition), clustered by gene expression and colored by cell type (left). UMAP plots showing cardiac cell type clusters across samples for the heart scRNA-seq data (right). **c)** 8,243 spatial transcriptomes of cardiac tissue sections from mock-infected and reovirus-infected mice at 4 and 7 dpi (one animal per condition). Hematoxylin and Eosin (H&E) stained image of reovirus-infected myocarditic tissue section used for spatial transcriptomics at 7 dpi (in box). **d)** UMAP plot of 7,695 single-cell transcriptomes from mock-infected and reovirus-infected ileum at day 1 and 4 dpi, clustered and colored by cell type (left). UMAP plots showing the gaussian kernel density of cells across samples for the ileum scRNA-seq data (right). **e)** Dot plot showing the percent of cells with non-zero viral transcripts and the mean viral transcript counts (UMIs) in ileal and cardiac cell types. **f)** RNA FISH labelling of cardiac cell type markers (*Tnnt2* for cardiomyocytes, *Postn* for fibroblasts, and *Cdh5* for endothelial cells), and immunofluorescence staining of reovirus antigen on a consecutive section showing infected endothelial cells within the infection foci at 7 dpi. Representative heart images from six biological replicates.

To faithfully identify reovirus transcripts in the ileum and heart, which are not polyadenylated, we performed hybridization-based enrichment of viral fragments captured in the scRNA-seq libraries (**Methods, Supp Fig. 4A-C**). In the ileum, we captured a total of 13,100 unique viral transcripts, with viral load decreasing from 1 dpi to 4 dpi. At 1 dpi, entero-endocrine cells had the highest fraction of infected cells followed by enterocytes and goblet cells, all of which are present in the gut epithelium. Lymphatic endothelial cells were infected at 4 dpi, suggesting that the virus reaches the bloodstream via lymphatic drainage to allow transmission of the virus to secondary sites in the body, including the heart, as shown before^12^ (**Supp Fig. 4D, Fig 1E**). We captured 2,762 unique viral transcripts from 392 cells in the T1L-infected hearts. The viral load first increased from 4 dpi to 7 dpi and then decreased from 7 dpi to 10 dpi, consistent with viral titer assays performed on whole hearts^9,13^ (**Fig. 1E, Supp Fig. 4E**). Endocardial and endothelial cells were the most frequently infected cell types at 4 dpi, suggesting that endocardial cells lining the ventricular lumen and endothelial cells lining the cardiac vasculature are among the first cells to be infected (**Fig. 1E**). We detected an increased infection in endothelial cells from 4 dpi to 7 dpi, consistent with viral titer assays performed on whole hearts^9,13^ (**Fig. 1E, Supp Fig. 4E**). We further detected viral transcripts in neutrophils, dendritic cells, and T cells in the 7 dpi heart (**Fig. 1E, Supp Fig. 4E)**. This observation suggests that antigen-presenting cells and immune cells may contribute to the spread of infection to other organs in the body. The role of infected dendritic cells in bringing more reovirus to the cardiac tissue during systemic infection has been discussed previously^8^.

To validate these observations, we performed histology, multiplexed RNA fluorescence in-situ hybridization (FISH), and immunofluorescence assays on tissue sections from myocarditic hearts and controls (multiple infected mice litters, **Supp Fig. 5A-5E, Methods**). We used RNA-FISH to visualize expression of genes specific to cardiomyocytes, fibroblasts, endothelial cells, macrophages, dendritic cells, neutrophils, and T cells (**Supp Fig. 5C-5E and Fig. 1F, Methods**). These experiments revealed infection foci and immune infiltration in myocarditic regions. We found *Itgam*+ *C1qa*-dendritic cells and *Trbc2*+ T cells inside the myocarditic regions and *S100a8*+ neutrophils in the border zones. In contrast, most *Itgam*+ *C1qa*+ macrophages were found outside the myocarditic regions at 7 dpi (**Supp Fig. 5C, 5D**). On consecutive tissue sections, we labeled reovirus antigen using immunofluorescence to identify reovirus infected cells (**Supp Fig. 5A, Fig. 1F**). Co-labelling for the endothelial cell marker *Cdh5* and reovirus transcript M3 on the same tissue sections confirmed the presence of viral transcripts in a subset of cardiac endothelial cells (**Supp Fig. 5E**). Endothelial cells that were positive for the reovirus antigen colocalized with T cells within the myocarditic regions (**Fig. 1F**). A small number of fibroblasts were often located on the edges of these regions (**Fig 1F**). Collectively, these results indicate that vascular endothelial cells are targets of reovirus in the heart.

### Endothelial cells are primed with a basal interferon response and play an important role in initiating host innate immune responses

To detect early transcriptional differences in the cardiac tissue after infection, we performed Differential Gene Expression Analysis (DGEA, mock vs infected hearts at 4 dpi, **Methods**). This analysis revealed a significant upregulation of 226 genes in the infected heart (two-sided Wilcoxon test, log fold-change > 1.0 and p-value < 0.01), including genes related to the interferon-β pathway, interferon signaling, and innate immune responses (**Supp Fig 6A-6B, Fig. 2A**).

**Figure 2:**
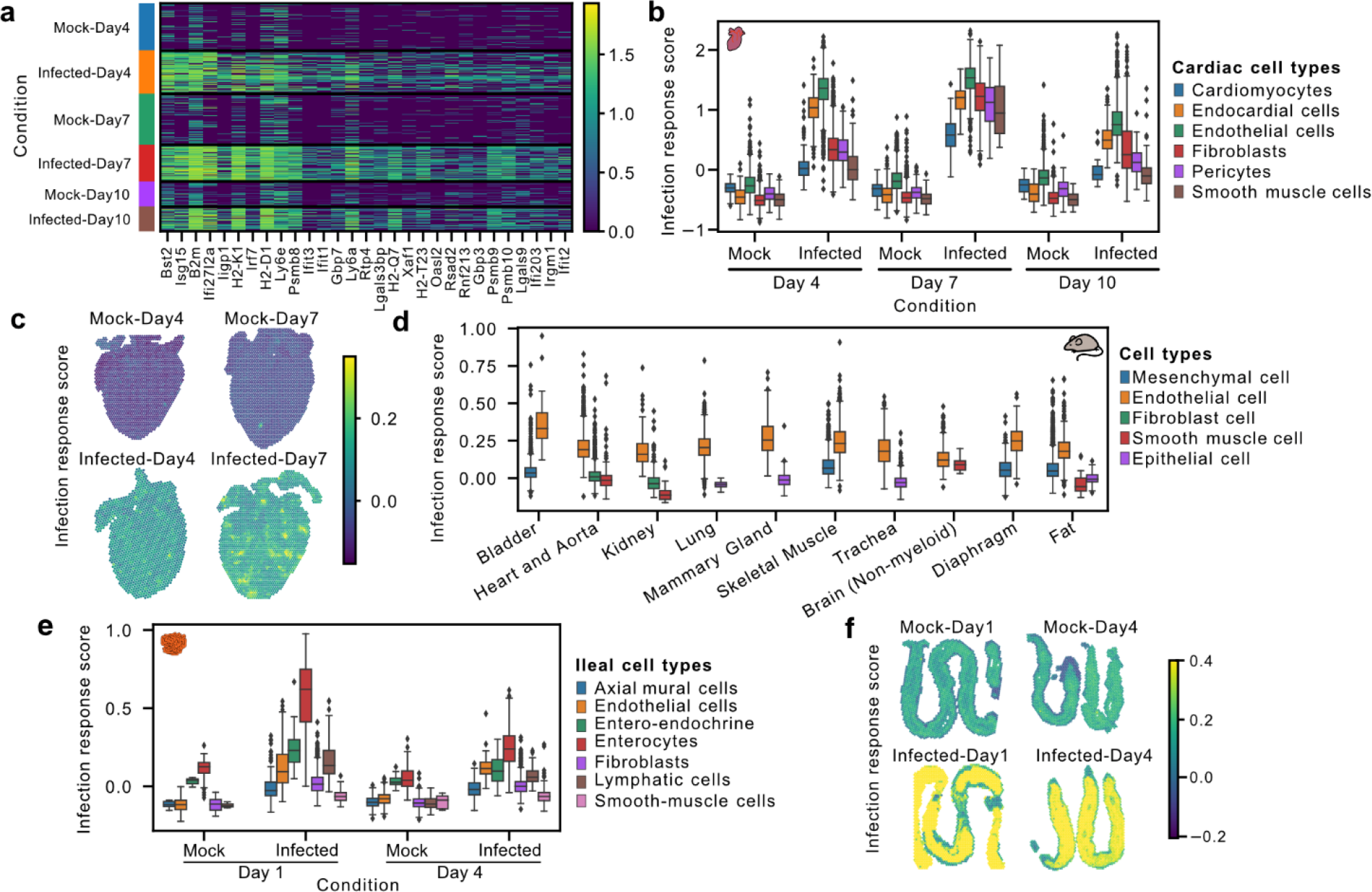
Endothelial cells have the highest basal interferon response and the highest increase in innate response upon reovirus infection. **a)** Heatmap showing the expression of the 25 most upregulated genes in the reovirus-infected heart as compared to mock at 4 dpi. **b)** Infection response score for cardiac cell types in scRNA-seq data across mock-infected and reovirus-infected hearts at three distinct stages. The infection response score represents the gene module score for a panel of 226 genes that are significantly upregulated (two-sided Wilcoxon test, log fold-change > 1.0 and p-value < 0.01) in the reovirus-infected heart as compared to the mock-infected heart at 4 dpi. **c)** Infection response score (defined above) across spots in spatial transcriptomics data. **d)** Infection response score calculated for five common cell types across 13 tissues from the tabula-muris mouse atlas data. **e)** Infection response score for ileal cell types in scRNA-seq data across mock-infected and reovirus-infected ileum at two distinct stages. The infection response score represents the gene module score for a panel of 438 genes significantly upregulated (two-sided Wilcoxon test, log fold-change > 1.0 and p-value < 0.01) in the reovirus-infected ileum at 1 dpi as compared to the mock-infected ileum. **f)** Infection response score for spatial transcriptomics data from mock-infected and reovirus-infected ileum at two distinct stages.

To quantify and compare the overall magnitude of early infection responses across different cell types, we computed a gene module score (infection response score, IR, module of 226 genes selected above). Comparison of the IR of different cell types in the absence of infection revealed a small, but higher IR in endothelial cells as compared to other cardiac cell types (**Fig. 2B**). In response to infection, an increase in IR was observed for all cardiac cell types, but the greatest increase in IR was observed for endothelial cells (**Fig. 2B**). These data suggest that endothelial cells lining the cardiac vasculature are important initiators of the host defense to viral infection. Comparison of IR scores using the spatial transcriptomic data showed increased IR scores in the infected hearts at 4 and 7 dpi with the highest scores found in myocarditic regions (**Fig. 2C**). Given our observation that endothelial cells within the heart had the highest IR score in the absence of infection, we asked if this observation was unique to heart tissue or was a more general phenomenon. To this end, we used the Tabula Muris scRNA-seq mouse atlas^14^ and estimated the IR of ∼16,000 cells of five major cell types (epithelial cells, fibroblasts, endothelial cells, smooth muscle cells, and mesenchymal cells) across 10 different organs and tissues. This analysis revealed that endothelial cells consistently had the highest IR score across all tissues in mice (**Fig. 2D**). These results indicate that endothelial cells lining the vasculature have a higher basal expression of innate response genes within most tissues, which may prime these cells to respond to viral dissemination within the blood and lymphatics.

To investigate the cell-type-specific IR in the ileum, the primary site of reovirus infection, we performed DGEA on reovirus-infected and mock-infected ileal cells at 1 dpi and found a significant upregulation of 438 genes (two-sided Wilcoxon test, log fold-change > 1.0 and p-value < 0.01), related to the interferon-beta pathway, interferon signaling, and innate immune responses in reovirus-infected ileal cells (**Supp Fig. 6C-6D**). We computed an IR score using this module of 438 genes and observed higher basal IR scores in enterocytes and entero-endocrine cells as compared to other ileal cell types (**Fig. 2E**). Enterocytes further showed the highest increase in IR score after infection, followed by entero-endocrine, endothelial, and lymphatic cells (**Fig. 2E**). Comparison of IR scores for spatial transcriptomic data further supported our analysis of the scRNA-seq data, showing increased IR scores in the infected ileum at 1 and 4 dpi with the highest scores evident within intestinal mucosa and villi (**Fig. 2F**). The intestinal epithelial cells must tolerate commensal microorganisms present in the lumen of the gut and yet still be responsive to invasive pathogens. Our data suggest that to achieve this, enterocytes and entero-endocrine cells in the gut epithelium are primed with a basal interferon response and play an important part in mounting innate immune responses in the early stages of viral infection.

### Inflamed endothelial cells recruit cytotoxic T cells and undergo pyroptotic cell death

To explore the heterogeneity of endothelial cell phenotypes in more detail, we reclustered all 9,786 cardiac endothelial cells in the scRNA-seq data. We observed four distinct phenotypes: ***i)*** uninflamed venous endothelial cells expressing *Nr2f2* and *Aplnr* mainly derived from the mock controls^15^, ***ii)*** arterial endothelial cells expressing *Gja4, Gja5*, and *Cxcl12* derived from both mock and infected cardiac hearts^15^, ***iii)*** inflamed endothelial cells derived from infected hearts at 4 and 10 dpi, and ***iv)*** inflamed endothelial cells from the heart at 7 dpi, with both inflamed endothelial cell clusters expressing *Isg15, Iigp1*, and *Ly6a* (**Fig. 3A-B**). DGEA across endothelial subclusters revealed that the inflamed 7 dpi endothelial cells overexpressed chemokines *Cxcl9* and *Cxcl10*, which are generally involved in immunoregulatory and inflammatory processes, but more specifically in the recruitment of T cells and NK T cells^16^ (**Fig. 3B-3C, Supp Fig. 7A**). In line with this observation, T cells in the 7 dpi hearts expressed the *Cxcr3* receptor (see below). The *Cxcl9*-high inflamed endothelial cells furthermore expressed high levels of cell adhesion marker genes *Vcam1* and *Icam1*, which help immune cells in the blood to attach to endothelial cells^17^ (**Fig. 3C, Supp Fig. 7A, 7E**). The endothelial cells also overexpressed MHC class 1 (*H2-D1* and *H2-K1*) and MHC class 2 (*Cd74*) molecules, suggesting their involvement in antigen presentation to adaptive immune cells (**Fig. 3B-C, Supp Fig. 7A, 7E)**. Endothelial cells have been shown to be involved in antigen presentation and shaping the cellular immune response in infectious myocarditis^17,18^. Gene ontology (GO) term enrichment analysis identified pathways further supporting the *Cxcl9*-high endothelial cells’ involvement in leukocyte cell-cell adhesion, T cell activation, regulation of interleukin-8 production, and response to cytokines, interferon-gamma, interleukin-1, and tumor necrosis factors (**Supp Fig. 7B**).

**Figure 3:**
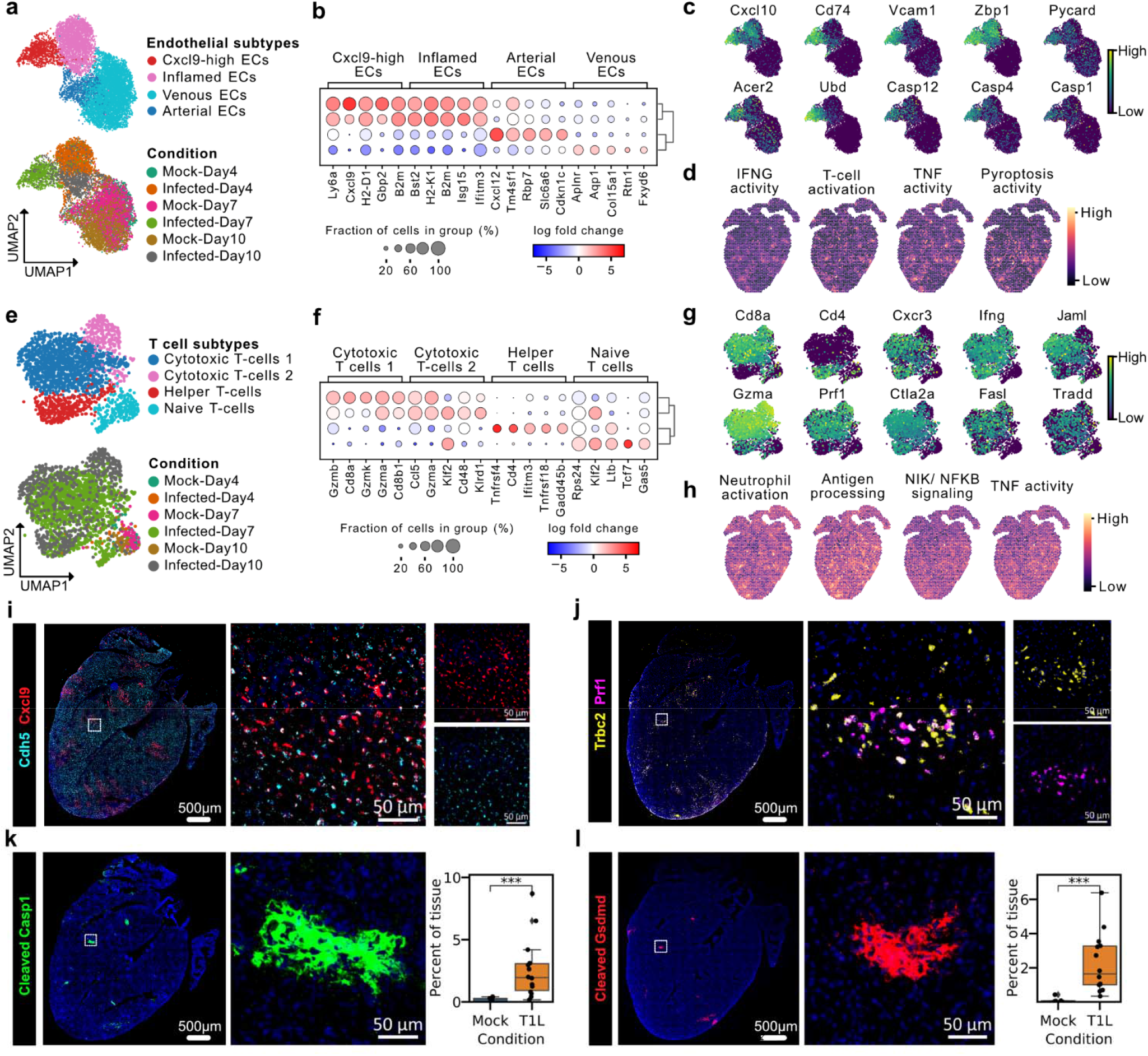
Cytotoxic T cells recruited by inflamed endothelial cells induce pyroptosis in myocarditic tissue. **a)** UMAP plot of 9,786 single-cell endothelial cell transcriptomes from mock-infected and reovirus-infected hearts at 4, 7, and 10 dpi colored by endothelial cell (EC) subtype clusters (phenotypes) (top) and condition (bottom). **b)** Heatmap showing top-five differentially expressed genes (two-sided Wilcoxon test, log fold-change > 1.0 and p-value < 0.01) for endothelial cell subtypes. **c)** UMAP plot showing the expression of genes upregulated in *Cxcl9*-high endothelial cells. **d)** Spatial transcriptomic maps of cardiac tissue from reovirus infected hearts at 7 dpi showing gene module scores calculated for four GO terms enriched in *Cxcl9*-high endothelial cells. **e)** UMAP plot of 2,205 single-cell T cell (TC) transcriptomes from mock-infected and reovirus-infected hearts at 4, 7, and 10 dpi colored by T cell subtype clusters (top) and condition (bottom). **f)** Heatmap showing top-five differentially expressed genes (two-sided Wilcoxon test, log fold-change > 1.0 and p-value < 0.01) for T cell subtypes. **g)** UMAP plot showing the expression of genes upregulated in cytotoxic T cells from myocarditic heart at 7 dpi. **h)** Spatial transcriptomics maps of cardiac tissue from reovirus infected hearts at 7 dpi showing gene module scores calculated for four GO terms enriched in cytotoxic T cells. **i**,**j)** RNA FISH staining for **i)** endothelial marker *Cdh5*, and chemokine *Cxcl9* **j)** T cell marker *Trbc2* and lytic molecule *Prf1* on consecutive sections from myocarditic hearts at 7 dpi. Representative images from 14 biological replicates (n=7 males and n=7 females). **k**,**l)** Immunofluorescence staining for **k)** cleaved Caspase1 protein-subunit (Casp1 p20 subunit) **l)** cleaved Gasdermin D protein (GSDMD N terminus fragment) on myocarditic hearts at 7 dpi. Representative images from 14 reovirus-infected biological replicates (n=7 males and n=7 females). Immunofluorescence signal from reovirus-infected hearts was compared to mock-infected hearts using two-sided Wilcoxon statistical test. p-value annotation legend: ns: p <= 1.00e+00, *: 1.00e-02 < p <= 5.00e-02, **: 1.00e-03 < p <= 1.00e-02, ***: 1.00e-04 < p <= 1.00e-03, ****: p <= 1.00e-04.

The observation that endothelial cells are involved in the recruitment of T cells prompted us to explore the heterogeneity of T cells in the infected hearts in more detail. To this end, we reclustered 2,205 T cell single-cell transcriptomes, leading to four subclusters representing three T cell subtypes, ***i)*** *Cd8+* cytotoxic T cells, ***ii)*** *Cd4+* helper T cells, and ***iii)*** naive T cells (**Fig 3E-F**). Both the cytotoxic and helper T cells identified within infected hearts expressed *Cxcr3* receptor, interferon-gamma (*Ifng)*, and the chemokines *Ccl3, Ccl4, Ccl5, S100A4*, and *S100A6*, suggesting their involvement in neutrophil recruitment and activation (**Fig. 3G, Supp Fig. 7C**). The *Cxcr3* receptor binds selectively to the chemokines *Cxcl9* and *Cxcl10*, promoting chemotaxis (**Fig. 3G**). Cytotoxic T cells represented the majority of infiltrating T cells and expressed *Prf1, Gzma, Gzmb*, and *Gzmk*, coding for lytic molecules associated with the granzyme-dependent exocytosis pathway^19^ (**Fig. 3F-G, Supp Fig. 7C, 7G**). These cells also expressed tumor necrosis factor superfamily genes *Fasl* and *Tradd*, which are involved in the Fas-induced cell death pathway. *Fasl* binds to *Fas* on the surface of target cells and mediates programmed cell death signaling and NF-κB activation (**Fig. 3G**). The Fasl-Fas apoptosis pathway is important in regulating T cells, in promoting tolerance to self-antigens, and is a mechanism by which cytotoxic T cells kill target cells^19^. GO term enrichment analysis identified pathways involved in neutrophil activation and degranulation, processing and presentation of exogenous peptide antigen, interleukin-1-mediated signaling pathway, tumor necrosis factor-mediated signaling, NF-κB-inducing kinase (NIK) /NF-κB signaling, cellular response to hypoxia, and apoptotic processes (**Supp Fig. 7D**).

The downstream gene markers for cell death*-*associated pathways *Pycard, Acer2, Zbp1*, and Caspases *Casp1, Casp4*, and *Casp12* were enriched in the *Cxcl9*-high endothelial cells, raising the possibility that cytotoxic lymphocytes are responsible for inflamed endothelial cell death (**Fig. 3B-3C, Supp Fig. 7E**). GO term enrichment of endothelial cells confirmed an upregulation of cell death pathways including activation of cysteine-type endopeptidase activity involved in the apoptotic process, positive regulation of the extrinsic apoptotic signaling pathway, and pyroptosis pathway (**Supp Fig. 7B**). We assessed the spatial transcriptomic data to validate direct interactions between *Cxcl9*-high inflamed endothelial cells and T cells and found that they were indeed spatially co-localized in the myocarditic regions and the border zone (**Supp Fig. 2C**). We calculated gene module scores for genes associated with ontology terms enriched in *Cxcl9*-high endothelial and cytotoxic T cells for spatial transcriptomics data and found these pathways to be enriched in the myocarditic regions (**Fig. 3D, 3H**, and **Supp Fig. 7E-H**).

We used histology, multiplexed RNA FISH, and immunofluorescence to validate our spatial transcriptomic and scRNA-seq findings on matched tissue sections from myocarditic and mock-infected hearts. (**Supp Fig. 8A, 8B, Methods**). The RNA FISH experiments confirmed the presence of Cxcl9-high endothelial cells (detected with *Cdh5*) colocalized with infiltrating T cells within myocarditic tissue (detected by Trbc*2*, and lytic molecule *Prf1*, **Supp Fig. 8A, 8B, Fig. 3I, 3J**). By immunofluorescence microscopy, we found expression of the pyroptosis-mediated cell death marker Caspase1 protein, the active cleaved Caspase1 protein and the pore-forming cleaved Gasdermin-D protein in myocarditic hearts at 7 dpi (consecutive tissue sections, **Supp Fig. 8C, Supp Fig. 8D, 8E** and **Fig. 3K, 3L**). These observations support the hypothesis that inflamed endothelial cells undergo pyroptosis in reovirus-infected myocarditic hearts.

Collectively, these results suggest that endothelial cells lining the cardiac vasculature act as a blood-heart barrier and play an important role in the recruitment and activation of the host adaptive immune system. These cells may be the target of both direct viral damage and immune-mediated damage during reovirus-induced myocarditis. Damage to the microvasculature within the heart may then cause loss of blood supply and be a factor in the subsequent death of cardiomyocytes independent of direct viral replication.

### Spatially restricted cell-type-specific gene expression in myocarditic tissue

The spatially restricted nature of myocarditis motivated us to explore the spatial heterogeneity of gene expression in reovirus-infected hearts. Our initial clustering of the spatial transcriptomic data revealed distinct transcriptional programs for myocarditic regions, the tissue bordering these myocarditic regions, and the rest of the ventricular tissue (**Fig. 1C, 4A, Supp Fig. 2A**). Differential spatial gene expression analysis for these regions revealed upregulation in the myocarditic regions of cell-type markers for infiltrating immune cells, (*Cd8a* and *Gzma* for T cells, *Nkg7* for NK cells, *S100a8* for neutrophils), markers of inflammation (*Cd52* and *Lyc62*, **Supp Fig. 9A-B**), and chemokines and cytokines (*Ccl5, Ccl2, Cxcl9*, and *Cxcl10*). Analysis of the corresponding scRNA-seq data showed that *Ccl5* is expressed by dendritic cells, *Ccl2* by fibroblasts, and *Cxcl9* and *Cxcl10* by endothelial cells. The receptor for *Ccl2, Ccr2*, is expressed in macrophages, indicating that fibroblasts use the *Ccl2-Ccr2* axis for macrophage recruitment during myocardial inflammation, as described recently^20,21^ (**Supp Fig. 9C**). Collectively these analyses suggest that chemokine-producing endothelial cells and cytokine-producing fibroblast cells contribute to the recruitment of immune cells to the myocarditic tissue.

**Figure 4:**
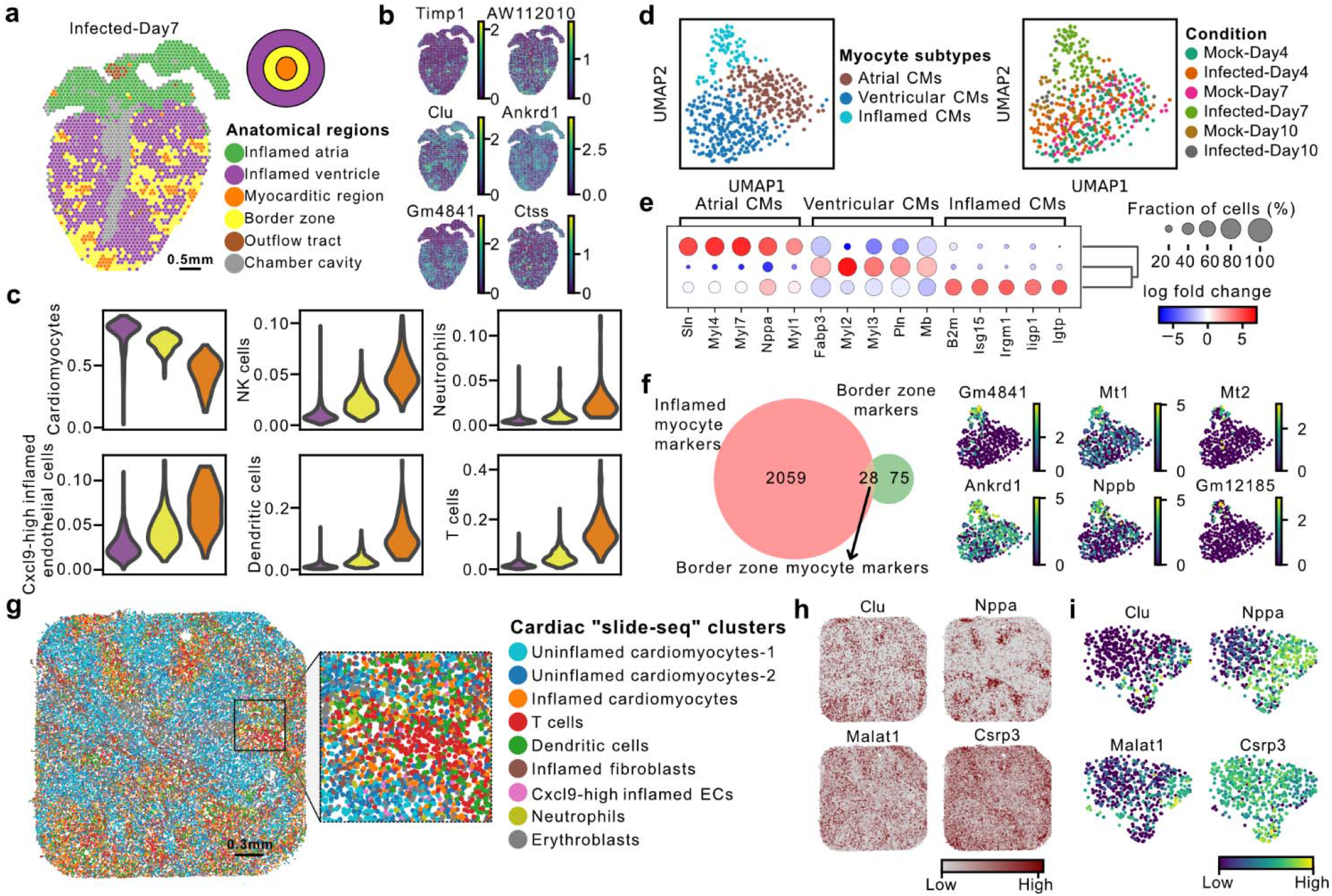
Myocarditic regions and the border zone have distinct transcriptomic profiles and cell type specific signatures. **a)** Spatial transcriptomics map of cardiac tissue section from reovirus-infected mice at 7 dpi colored by spot clusters representing transcriptionally distinct tissue regions. **b)** Spatial transcriptomics maps of cardiac tissue sections from reovirus-infected mice at 7 dpi showing the expression of differentially expressed genes of interest in the myocarditic and the border zone. **c)** Changes in average predicted cell-type proportions across the infected ventricle, for cell types enriched in the myocarditic region and the border zone. **d)** UMAP plot of 502 single-cell cardiomyocyte cell transcriptomes from mock-infected and reovirus-infected hearts at 4, 7, and 10 dpi colored by myocyte cell subtype (phenotypes) (left) and condition (right). **e)** Heatmap showing top-five differentially expressed genes (two-sided Wilcoxon test, log fold-change > 1.0 and p-value < 0.01) for cardiomyocyte cell subtypes. **f)** Venn Diagram showing myocyte-specific genes upregulated in the border zone around the myocarditic regions (left). UMAP plot showing the expression of myocyte-specific genes which are upregulated in the border zone of myocarditic regions (right). **g)** High-resolution Slide-seq spatial transcriptomics map of cardiac ventricular tissue from reovirus infected mice at 7 dpi colored by Slide-seq bead clusters. Zoom-in shows the spatial arrangement of Slide-seq clusters within a myocarditic region. **h)** Spatial transcriptomic maps showing Slide-seq expression of four cardiomyocyte specific genes enriched in inflamed cardiomyocytes as compared to uninflamed myocytes. **i)** UMAP plot showing the scRNAseq expression of myocyte-specific genes which are upregulated in inflamed myocytes in the slide-seq data.

Closer inspection of the myocarditic regions and border zones showed an upregulation of additional genes of interest, including *Timp1, AW112010, Clu, Ankrd1, Gm4841, and Ctss* (**Fig. 4B**). *Timp1* was mainly expressed by inflamed fibroblasts in the scRNA-seq data (**Supp Fig. 9D**). *Timp1* is a natural inhibitor of the matrix metalloproteinases (MMPs), a group of peptidases involved in the degradation of the extracellular matrix. Upregulation of *Timp1* in patients with deteriorating heart failure was reported previously^22^. *AW112010* was expressed by inflamed endothelial cells and fibroblasts in the scRNA-seq data. *AW112010* encodes an interferon-induced small secreted protein which regulates inflammation by suppressing IL-10 within proinflammatory T-cells^23^ (**Supp Fig. 9D**). *Clu* was expressed in a subset of inflamed cells from all cardiac cell types in our data. *Clu* is upregulated during severe myocarditis^24^ (**Supp Fig. 9D**). *Ctss* was expressed mainly in monocytes (**Supp Fig. 9D**). *Ctss* encodes a protease used for degradation of antigenic proteins to peptides for presentation by MHC class II molecules. Increased formation of immunoproteasomes in susceptible mice has been shown to affect the generation of antigenic peptides and subsequent T cell activity in viral myocarditis^25,26^. GO term analysis of genes upregulated in the border zone revealed enrichment of terms related to the response to tumor necrosis factor, response to interleukin-1, and NIK/ NF-κB signaling (**Supp Fig. 9E**).

To further understand the effect of immune cell infiltration on the cell type composition surrounding the myocarditic regions, we assessed cell type proportions as a function of distance from myocarditic regions in the tissue. We quantified the cell type proportions in myocarditic regions, the border zones, and the rest of the ventricular tissue, and found that the fraction of *Cxcl9*-high endothelial cells, *Ccl2*+ fibroblasts, T cells, dendritic cells, and NK cells was increased in the myocarditic regions, and the fraction of cardiomyocytes was reduced in myocarditic regions (**Fig. 4C, Supp Fig. 2C**). To understand the phenotype of *Ccl2*+ fibroblasts enriched in myocarditic region and border zone, we reclustered 9,192 fibroblast cells from the scRNA-seq dataset and identified a distinct cluster of inflamed *Ccl2*+ fibroblasts from the infected heart at 7 dpi (**Supp. Fig. 9F, 9G**). The *Ccl2*+ fibroblasts expressed high levels of MHC class 1 (*H2-D1* and *H2-K1*), adhesion marker genes *Vcam1* and *Icam1*, and other genes such as *Serpina3g, C3*, and *Ms4a4d* (**Supp Fig. 9H, 9I**). Moreover, these cells also expressed *Casp1* and *Casp4*, suggesting that fibroblasts also undergo pyroptosis (**Supp Fig. 9H**).

To investigate the effect of inflammation on cardiomyocytes in myocarditic hearts, we reclustered 502 cardiomyocytes from the scRNA-seq dataset and identified three distinct phenotypes: ***i)*** ventricular myocytes expressing *Myl2, Myl3*, and *Mb* derived from mock and infected hearts at 4 and 10 dpi, ***ii)*** atrial myocytes expressing markers *Myl4, Myl7*, and *Nppa* derived from mock and infected hearts at 4 and 10 dpi, and ***iii)*** inflamed myocytes from the infected heart at 7 dpi expressing innate immunity genes *Isg15, Igtp*, and *Iigp1*^27^ (**Fig. 4D-E**). Inflamed myocytes from the infected heart at 7 dpi had a distinct phenotype when compared to the myocytes from hearts at 4 and 10 dpi, which clustered with myocyte cells from mock-infected hearts (**Fig. 4E**). To find transcriptional signatures for myocytes present in the border zone, we selected genes that were both enriched in cardiomyocytes in the scRNA-seq data and upregulated in the border zone. This analysis revealed that cardiomyocytes in the border zone expressed *Gm4841, Gm12185, Mt1, Mt2, Ankrd1*, and *Nppb* (**Fig. 4F, Supp Fig. 9J**). *Gm4841* and *Gm12185* are interferon-inducible genes produced in response to interferon-gamma. *Mt1* and *Mt2* genes modulate inflammation and support remodeling in ischemic cardiomyopathy in mice^28^. Upregulation of *Ankrd1*, a myocyte survival factor, occurs during late-stage heart disease in patients with idiopathic dilated cardiomyopathy^29^. A recent study shows that cardiomyocytes expressing *Ankrd1* are localized in the border zone on day 1 post-myocardial infarction^30^.

To visualize the spatial distribution and phenotypes of cardiac cell types at higher spatial resolution, we also performed “Slide-seq” spatial transcriptomics^31,32^ (resolution = 10 μm) on ventricular tissue from a single reovirus-infected myocarditic heart **(Methods, Supp Fig. 10A)**. We performed unsupervised clustering and DGEA to label these near single-cell resolution slide-seq spatial transcriptomes as cardiac cell types (**Supp Fig. 10B, 10C** and **Fig. 4G**). We visualized the cell types on the spatial maps and performed neighborhood enrichment analysis and observed neutrophils, *Cxcl9-*expressing endothelial cells, and inflamed cardiomyocytes organized in close proximity to infiltrating T cells and dendritic cells in the myocarditic regions (**Supp Fig. 10D** and **10E**). We furthermore used deconvolution using the scRNAseq data as a reference to obtain cell type predictions and to quantify cell-type-specific gene expression at every spatial location (**Methods, Supp Fig. 11**). We compared the phenotypes of inflamed and uninflamed myocyte clusters using DGEA and confirmed the upregulation of *Ankrd1, Nppb, Gm4241*, and *Saa3*. We furthermore identified additional inflammation and stress related markers for inflamed cardiomyocytes such as *Clu*^33^, *Malat1*^34^, *Nppa*^35,36^, and *Cspr3*^37^ (**Supp Fig. 10F & 10G and Fig. 4H & 4I**). Together, our analysis reveals that tissue injury is localized to myocarditic regions with remodeling and stress programs being active in the border zone and demonstrates the importance of spatially resolved molecular measurements to study viral myocarditis.

### Reduced adaptive immune cell infiltration associated with reovirus K287T mutant

We recently reported a reovirus mutant T1L S4-K287T (K287T) which has a point mutation in the S4 gene encoding outer capsid protein sigma-3 (σ3), a double-stranded (ds) RNA-binding multifunctional protein that promotes viral protein synthesis and facilitates viral entry and assembly^9^. This mutation abolishes the capacity of σ3 to block dsRNA-mediated activation of protein kinase R (PKR). The T1L K287T mutant is less virulent than the WT strain in neonatal mice. K287T replicates to WT titers in the heart at 4 dpi, but to significantly lower viral titers than WT virus at 7 dpi. The K287T mutant does not induce myocarditis as observed by calcium staining in the tissue^9^. To confirm our findings about immune-mediated pathogenesis during reovirus infection, we performed additional scRNA-seq for K287T infected hearts at 4, 7, and 10 dpi. We generated a total of 16,771 single-cell transcriptomes and integrated the data with the data from the WT virus. We did not observe sample-specific clusters after data integration, suggesting minimal experimental batch effects (**Fig. 5A, Supp Fig. 12A**). We performed viral transcript enrichment and compared the mean viral transcripts in WT- and mutant-infected cells. We found similar levels of mean viral transcripts for WT and K287T viruses at 4 dpi but a 60-fold lower viral load for K287T at 7 dpi, consistent with viral titer assays^9^ (**Supp Fig. 12B-E**). We then compared the early cardiac cell type host responses to K287T and WT infection. K287T induced a similar level of innate immune responses as WT virus with endothelial cells showing the highest increase in cardiac IR score (as defined before) at 4 dpi (**Fig. 5B**).

**Figure 5:**
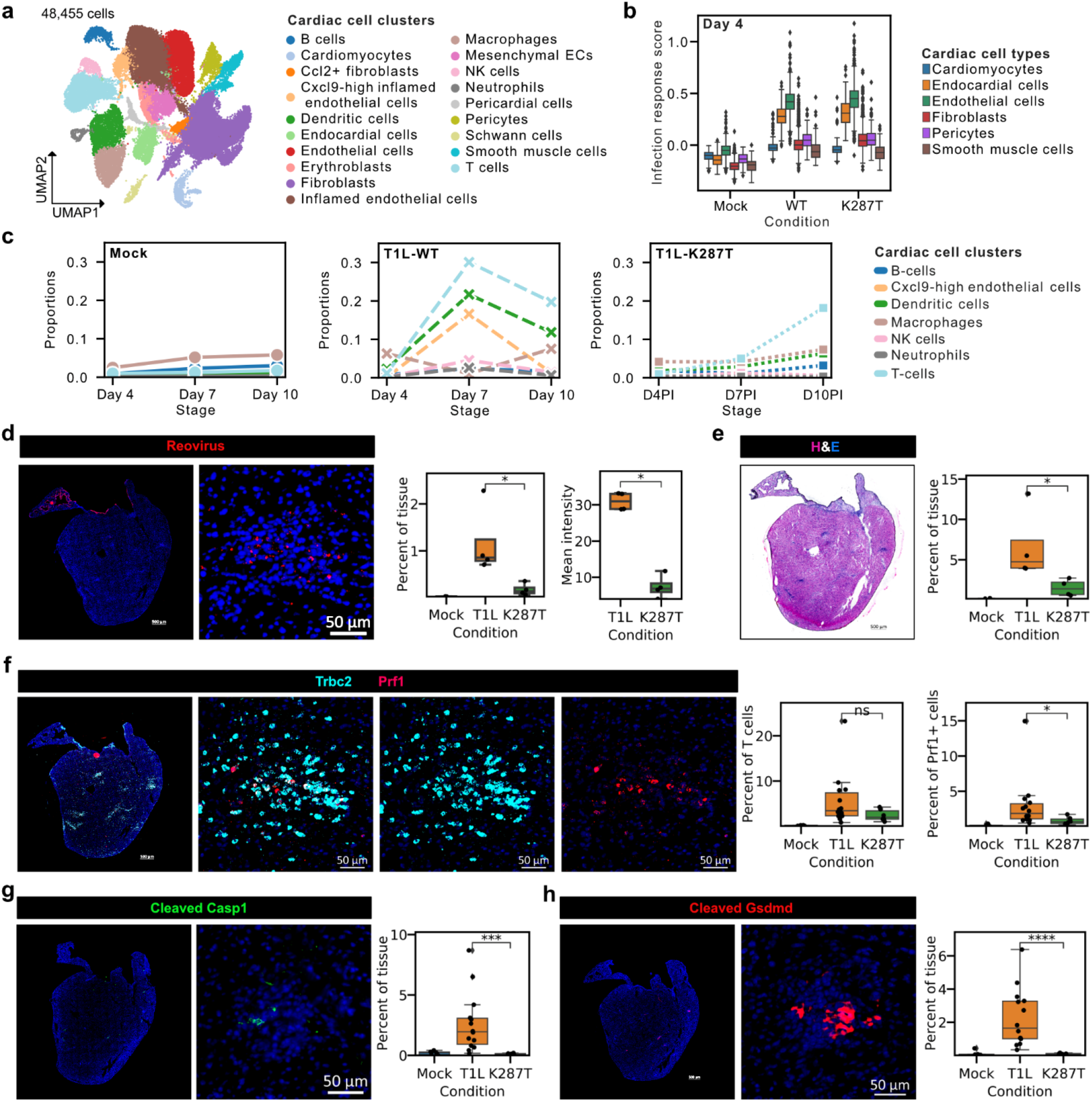
A robust innate immune response but reduced adaptive immune cell infiltration explains the non-myocarditic phenotype on infection with reovirus K287T mutant. **a)** UMAP plot of 48,455 single-cell cell transcriptomes from mock-infected, reovirus-wildtype (WT) infected, and reovirus mutant (K287T) infected hearts at 4, 7, and 10 dpi (one animal per condition) colored by cell-type clusters. **b)** Infection response score for cardiac cell types in scRNA-seq data across mock-infected, reovirus-WT infected, and reovirus-K287T infected hearts on 4 dpi. The infection response score represents the gene module score for a panel of 226 genes that are significantly upregulated in the reovirus-WT infected sample as compared to the mock-infected sample at 4 dpi. **c)** Changes in cell-type proportions with time for cell types detected in the myocarditic regions. Panels show the changes in cell-type proportions across mock-infected, reovirus-WT infected, and reovirus-K287T infected cells. **d)** Immunofluorescence images of reovirus antigen on reovirus mutant (K287T) infected hearts at 7 dpi. e) Hematoxylin and Eosin (H&E) stained image of K287T-infected heart tissue section at 7 dpi. **f)** RNA FISH staining for of T cell marker *Trbc2*, and lytic molecule *Prf1* on K287T-infected heart tissue section at 7 dpi. **d-f)**. Representative heart images from six K287T-infected hearts. **g-h)** Immunofluorescence staining for **g)** cleaved Caspase1 protein-subunit (Casp1 p20 subunit) **l)** cleaved Gasdermin D protein (GSDMD N terminus fragment) on K287T-infected heart tissue section at 7 dpi. Representative images from six K287T-infected biological replicates (n=3 males and n=3 females). Immunofluorescence signal from K287T-infected hearts was compared to WT-infected hearts using two-sided Wilcoxon statistical test. p-value annotation legend: ns: p <= 1.00e+00, *: 1.00e-02 < p <= 5.00e-02, **: 1.00e-03 < p <= 1.00e-02, ***: 1.00e-04 < p <= 1.00e-03, ****: p <= 1.00e-04.

We analyzed the cell type composition of inflamed *Cxcl9*-high endothelial cells and immune cells detected in K287T- and WT-infected hearts. We observed fewer *Cxcl9*-high endothelial cells and immune cells including cytotoxic T cells, infiltrating the heart at 7 dpi compared to WT-infected heart (**Fig. 5C**). These differences correlate with the reduced levels of inflammation associated with the K287T mutant **(Fig. 5E)**. To validate these observations, we performed RNA FISH and immunofluorescence staining on K287T-infected hearts and compared them to mock-infected and reovirus WT-infected hearts (**Fig. 5D-H**). Immunostaining for reoviral antigen in tissue sections confirmed both a significantly reduced area with viral replication (two-sided Mann-Whitney test, p-value < 0.05) and significantly lower viral antigen within those areas (two-sided Mann-Whitney test, p-value < 0.05), consistent with the scRNA-seq analysis and viral titer assays (**Supp. Fig. 12C, Fig. 5D**). We observed a reduction in infiltration of T cells in K287T-infected hearts as compared to WT-infected hearts at 7 dpi (**Fig. 5F**). The fraction of total cytotoxic immune cells (*Prf1*+) was significantly reduced in K287T-infected hearts as compared to WT-infected hearts (two-sided Mann-Whitney test, p-value < 0.05, **Fig 5F**). These findings support the reduced immune-mediated cytotoxicity seen in K287T-infected hearts. This was further supported by a significant reduction in cleaved Caspase1 and cleaved Gasdermin-D protein expression in K287T-infected hearts as compared to WT-infected hearts (two-sided Mann-Whitney test, p-value < 1.00e-03, **Fig. 5G, 5H**). Our results show that cardiac endothelial cells mount a potent and robust innate immune response when infected with the K287T mutant virus. Clearance of the K287T virus from most infected cells by 7 dpi leads to a lower immune-mediated cytotoxicity, which correlates with lack of cardiac injury. These results suggest that a robust early innate immune response in endothelial cells is critical for early viral clearance and prevention of subsequent cardiac injury mediated by cytotoxic immune cells during reovirus-induced myocarditis.

## DISCUSSION

Viral myocarditis has been recognized as a cause of heart failure for more than 50 years, but it is still a challenging disease to study, diagnose, and treat^38^. Here, we used integrated spatial and single-cell RNA-seq to dissect the temporal, spatial, and cellular heterogeneity of reovirus-induced acute myocarditis in a neonatal mouse model. We assayed ileum and heart tissues at multiple time points after infection. We investigated the cell types that are infected, and the cellular and spatial heterogeneity of innate and adaptive immune responses. We generated a total of thirteen scRNA-seq and eight spatial transcriptomics datasets, spanning two organs, four time points, and three infection conditions. Our data provide detailed insight into the chronology of molecular events that lead to reovirus-induced myocarditis. After oral inoculation, reovirus T1L infects entero-endocrine and enterocyte cells in the gut epithelium within 1 dpi. These cells mount a potent innate immune response to inhibit viral replication. The virus then infects the gut lymphatic cells within 4 dpi and is transmitted via lymphatics to the bloodstream and then to secondary sites in the body, including the heart. Around 4 dpi, the virus infects the endothelial cells lining the cardiac vasculature. Endothelial cells mount a potent innate immune response in the heart. In symptomatic cases, inflamed endothelial cells secrete chemokines that recruit circulating immune cells, including cytotoxic T cells. These inflamed endothelial cells then undergo pyroptotic cell death in the myocarditic tissue. Overall, our experiments reveal a dynamic and spatially heterogeneous network of cellular phenotypes and cell-cell interactions associated with reovirus-induced myocarditis.

Integrated high-throughput scRNA-seq and spatial transcriptomics was recently used to study heart development^39,40^ and heart disease^30,41^, but these methods have not been used to study viral myocarditis prior to our work. Bulk RNA-seq has been used previously to profile transcriptomic signatures of infection, inflammation, and tissue injury associated with viral myocarditis^9,42–45^. Yet, these ensemble-level approaches do not capture the cellular and spatial heterogeneity of host response to infection. scRNA-seq has recently been used to study Coxsackievirus B3 (CVB3)-induced myocarditis in a mouse model^46^. Lasrado et al. report inflammatory phenotypes of myeloid cells, the role of fibroblasts in remodeling and inflammation, and the role of cytotoxic T-cells in CVB3-induced myocarditis. However, the cardiac cell types that are targeted by the virus, the cell type heterogeneity in basal interferon response and innate immune response, and the spatial restriction of transcriptional programs were not explored in this study.

Reovirus infection occurs often in humans, but most cases are mild or subclinical. These viruses display a broad host range, but only young hosts develop the disease. After infection of neonatal mice, reoviruses cause injury to a variety of organs, including the heart, liver, and the central nervous system, depending on the viral strain. Reovirus Type-1-Lang (T1L) strain is mildly virulent and causes myocarditis in ∼50% of the infected mice. Neonatal mice with myocarditic hearts due to T1L infection survive with tissue damage and have an increased rate of heart failure. Therefore, reovirus T1L infection in neonatal mice is an ideal model to study the mechanisms and pathogenesis of reovirus induced myocarditis in young hosts. Previous studies have claimed that the direct cytopathic effect of viral replication on cardiac cells is the main cause of cardiac damage during reovirus-induced myocarditis^7,47^. Notably, Sherry et al. found that reovirus infection can induce myocarditis in immunodeficient mice lacking B and/or T cells, suggesting that reovirus-induced myocarditis does not strictly require adaptive immunity^7,11^. However, these previous experiments do not rule out the possibility that the host adaptive immune response can augment or delimit the nature and amount of host damage in immune-competent mice, as is suggested by our work. In addition, the viral strain used in these experiments was substantially more virulent. Holm et al. and Stewart et al. have studied the protective role of innate immune responses in reovirus-induced myocarditis^13,48^. However, prior to this study the temporal, spatial, and cell type heterogeneity of basal type-I IFN and innate immune responses to infection had not been characterized. Miyamoto et al. and Stewart et al. compared basal levels of type-I IFN between cardiac myocytes and fibroblasts *in vitro* but these studies did not include all the cell types that make up complex cardiac tissues^27,49^.

Spatiotemporal characterization of viral myocarditis is crucial to understanding the viral and host factors that are important for disease pathology. This knowledge may ultimately lead to novel diagnostic approaches and better treatments. Several viruses that frequently infect humans can cause myocarditis, including Adenovirus, enteroviruses, Epstein-Barr virus, human Herpesvirus 6, parvovirus B19, and SARS-CoV2. The results presented here may not be representative of the mechanisms for other viral causes of myocarditis or viral myocarditis in adult hosts. However, the approaches that we have implemented here can be used in future studies to investigate how the induction, pathophysiology, and course of myocarditis induced by these viruses differs. We hope that the data and analysis routines that we make available here will be a valuable resource for such future studies.

## METHODS

### Reovirus infections of neonatal C57BL/6J mice

Confirmed pregnant female C57BL/6J mice were ordered from Jackson Laboratories to be delivered at embryonic stage E14.5. Litters weighing 3 gram/ pup were gavaged using intramedic tubing (Becton Dickinson 427401) per os with 50 μl with 10^7^ PFU reovirus type 1 lang (T1L): wildtype or K287T mutant in 1x phosphate buffered saline (PBS) containing green food color (McCormick) via a 1ml tuberculin slip tip syringe (BD 309659) and 30G x 1/2 needle (BD 305106). Litters treated with 1x PBS containing green food color alone on the same day were used as mock controls for the respective infection groups. The mock-infected and reovirus-infected mice pups were weighed daily until the time points used in the study (1-, 4-, 7-, and 10-days post infection (dpi)). Due to the difficulty in determining the sex of mice during infection and early neonatal stages, we randomly selected the mice to collect ileum and heart tissues for scRNAseq and spatial transcriptomics experiments (**Supp Data 2**). All animal work was conducted ethically, conforming to the U.S. Public Health Service policy, and was approved by the Institutional Animal Care and Use Committee at Cornell University (IACUC Number 2019-0129).

### Sample preparation for single-cell transcriptomics of cardiac tissue

We sacrificed mock-infected and reovirus-infected C57BL/6J mice on day 4, day 7, and day 10 post-infection and collected cardiac tissues for single cell transcriptomics. Single heart tissue from respective stages (one heart per stage) were isolated aseptically, washed with ice-cold Hank’s Balanced Salt Solution, HBSS (with calcium and magnesium chloride; Gibco 14025-134), and minced into 1-2mm pieces. Cardiac tissue pieces were then digested in tissue dissociation media with 200U/mL collagenase type II (Gibco 17100-015), 1 mg/ml dispase (Sigma D4693), and 3mM calcium chloride in HBSS for four cycles of 10 minutes under mild agitation at 37°C in 1.5 ml eppendorf tubes. After every 10-minute cycle, cell suspension was collected, added to ice-cold 1x PBS with 0.04% bovine serum albumin (BSA; Sigma A3803) and new dissociation media was added to the tubes. At the end of the digestion, the cells were passed through a 70µm filter and centrifuged into a pellet. To remove most blood contaminants, samples were resuspended in an ammonium-chloride-potassium (ACK) lysis buffer (Lonza #10-548E) for 3-5 minutes and centrifuged. Samples were then washed again in PBS with 0.04% BSA and then resuspended at 1□×□10^6^ cells per ml. Cells from each sample were stained with Trypan Blue and cell viability was calculated on an automated cell counter (Countess II) before loading the cells on 10x Chromium. We used these cell viabilities to adjust the number of cells loaded on 10x Chromium to get the desired number of transcriptomes from viable cells for each sample (5000 cells per sample).

### Sample preparation for single-cell transcriptomics of intestinal tissue

We sacrificed mock-infected and reovirus-infected C57BL/6J mice on days 1 and 4 post-infection and collected intestinal ileum tissue for single cell transcriptomics. Single intestinal ileum tissue from respective stages (one tissue per stage) were isolated aseptically, washed with ice-cold Hank’s Balanced Salt Solution, HBSS (without calcium and magnesium chloride; Gibco 14175-095) to remove contamination. The ileum tissue was then opened longitudinally, washed again with HBSS, and minced into 1-2mm pieces. To isolate the epithelial layer of cells, ileum tissue pieces were incubated in HBSS with 10mM Ethylenediaminetetraacetic acid (EDTA, Invitrogen 15575-038) and 1mM Dithiothreitol, (DTT, Sigma 43816-10ML) for two cycles of 10 minutes under mild agitation at 37°C. After every 10-minute cycle, cell suspension containing the intestinal epithelial cells was collected, added to ice-cold 1x PBS with 0.04% bovine serum albumin (BSA; Sigma A3803). The undigested pieces of lamina propria were then washed thoroughly with PBS (with calcium and magnesium chloride; Gibco 14080-055) to get rid of all EDTA. These pieces were then transferred to fresh tubes and incubated in 200U/ml Collagenase type I (Gibco 17100-017) and 3mM calcium chloride in PBS for three cycles of 10 minutes under mild agitation at 37°C in 1.5 ml eppendorf tubes. After every 10-minute cycle, cell suspension containing the lamina propria cells was collected, added to ice-cold phosphate buffered saline, PBS with 0.04% BSA in separate tubes. At the end of the digestion, the cells were passed through a 40µm filter and washed twice in PBS with 0.04% BSA and then resuspended at 1□×□10^6^ cells per ml. Cells from intestinal epithelium and lamina propria for each sample were stained with Trypan Blue and cell viability was calculated on automated cell counters (Countess II). Cell counts adjusted with viability were then pooled as 40% epithelial cells and 60% lamina propria to adjust the number of cells loaded on 10x Chromium and to get the desired number of transcriptomes from viable cells for each sample (5000 cells per sample).

### Single-cell RNA sequencing library preparation

5000-6000 viable cells per sample (for heart and ileum tissues) were targeted on the Chromium platform (10x Genomics) using one lane per sample per time point. Single-cell libraries were built using the Chromium Next GEM Single Cell 3’ Library Construction V3 Kit (10x Genomics) and were then sequenced on an Illumina NextSeq 500 using 75 cycle high output kits (Index 1 = 8, Read 1 = 28, and Read 2□=□55) for all samples. Sequencing data were aligned to a combined mouse and reovirus reference genome (described below) using the Cell Ranger 6.0.0 pipeline (10x Genomics).

### Hybridization-based enrichment of viral fragments

We performed a hybridization-based enrichment of viral fragments on a part of scRNA-seq libraries using xGen NGS target enrichment kit (IDT; 1080577). In this approach, a panel of 5’-biotinylated oligonucleotides is used for capture and pulldown of target molecules of interest, which are then PCR amplified and sequenced. We designed a panel of 202 biotinylated probes tiled across the entire reovirus T1L genome to selectively sequence viral molecules from the scRNA-seq libraries **(Supp Data 3)**. 300ng of fragmented and indexed scRNA-seq libraries from reovirus-WT infected hearts, reovirus-mutant infected hearts, and reovirus-infected ileum were pooled in three separate reactions for xGen hybridization capture. Two rounds of hybridization capture using the xGen enrichment protocol were performed for every reaction to enrich viral transcripts. Amplification was performed for a total of 18 PCR cycles after the first round of capture. 50% of the amplified product was used for the second round of hybridization capture and amplification was performed for a total of 5 PCR cycles after the second round of enrichment. Post-enrichment products were pooled and sequenced on Illumina Mini-seq for ileum libraries and NextSeq 500 for heart libraries.

### Sample preparation for Visium spatial transcriptomics

Whole hearts and intestinal ileum were isolated using aseptic techniques and placed in ice cold sterile Hank’s Balanced Salt Solution, HBSS (without calcium and magnesium chloride; Gibco 14175-095). Blood and other contamination were carefully removed by perfusing the tissues with fresh HBSS. Fresh tissues were immediately embedded in Optimal Cutting Compound (OCT) media (SAKURA 25608-930) and frozen in a liquid-nitrogen-cooled isopentane (EMD Millipore, MX0760) bath for spatial transcriptomics experiments. The tissue blocks were cut into 10µm sections using Thermo Scientific CryoStar NX50 cryostat and mounted on Visium Gene Expression slides (10x Genomics), which were pre-cooled to -20°C and used for the Visium Spatial Gene Expression experiment.

### Visium spatial transcriptomics library preparation

We used the Visium Spatial Gene Expression (10x Genomics) platform for the spatial transcriptomics experiments. Tissue sections from fresh-frozen hearts (mock-infected and reovirus-infected at day 4 and day 7 post infection) and ileum (mock infected and reovirus infected at day 1 and day 4 post infection) were mounted with one section per capture area on individual Visium Gene Expression slides. These sections are then fixed in pre-chilled methanol for 30 minutes and then hematoxylin and eosin (H&E) stained and imaged, which is later used by the 10x Genomics Space Ranger (version 1.0.0) software to detect the spots which are covered by the tissue. The optimal permeabilization time for 10 µm thick sections was found to be 18 minutes for the heart and 12 minutes for the ileum using the 10x Genomics Visium Tissue Optimization kit. Spatially tagged cDNA libraries were built using the 10x Genomics Visium Spatial Gene Expression 3’ Library Construction V1 Kit. H&E-stained heart tissue sections were imaged using Zeiss PALM MicroBeam laser capture microdissection system at 20x objective and the images were stitched and processed using Fiji ImageJ software. cDNA libraries were sequenced on an Illumina NextSeq 500/550 using 150 cycle high output kits (Read 1□= 28, Read 2□=□120, Index 1□=□10, and Index 2□=□10) for ileum and on an Illumina NextSeq 2K (P2 flow cell) using the 100-cycle kit (Read 1 = 28, Read 2 = 96, Index 1□=□10, and Index 2□=□10) for the heart samples. Fiducial frames around the capture area on the Visium slide were aligned manually and spots covering the tissue were selected using Loupe Browser 4.0.0 software (10x Genomics). Sequencing data was then aligned to a combined mouse and reovirus reference genome (described below) using the Space Ranger 1.0.0 (10x Genomics) pipeline to derive a feature spot-barcode expression matrix. Visium slide number V19B23-046 was used for spatial transcriptomics experiment on mice hearts (mock-infected 4 dpi: capture area D1, reovirus-infected 4 dpi: capture area B1, mock-infected 7 dpi: capture area C1, and reovirus-infected 7 dpi: capture area A1). Visium slide number V19B23-045 was used for spatial transcriptomics experiment on mice ileum tissue (mock-infected 1 dpi: capture area D1, reovirus-infected 1 dpi: capture area B1, mock-infected 4 dpi: capture area C1, and reovirus-infected 4 dpi: capture area A1).

### Sample preparation for Slide-seq spatial transcriptomics

Whole hearts were isolated using aseptic technique and placed in ice cold sterile Hank’s Balanced Salt Solution, HBSS (without calcium and magnesium chloride; Gibco 14175-095). Blood and other contamination were carefully removed by perfusing the tissues with fresh HBSS. Fresh tissues were immediately embedded in Optimal Cutting Compound (OCT) media (SAKURA 25608-930) and frozen in a liquid-nitrogen-cooled isopentane (EMD Millipore, MX0760) bath for spatial transcriptomics experiments. The tissue blocks were cut into 10µm sections using Thermo Scientific CryoStar NX50 cryostat and mounted on a “Curio Seeker Tile (Tile ID #A0004_043, Curio Bioscience). A barcode whitelist and a barcode position file for the corresponding tile were provided by Curio Bioscience.

### Slide-seq spatial transcriptomics library preparation

Slide-seq spatial transcriptomics experiment was performed using the Curio Seeker Kit (Curio Bioscience) according to manufacturer’s instructions. Briefly, a tissue section from a fresh-frozen reovirus-infected heart at 7 dpi was mounted on a 3mmx3mm spatially indexed bead surface (Curio Seeker Kit, Tile ID #A0004_043, Curio Bioscience). After RNA hybridization and reverse transcription, the tissue section was digested, and the beads were removed from the glass tile and resuspended. Second strand synthesis was then performed by semi-random priming followed by cDNA amplification. A sequencing library was then prepared using the Nextera XT DNA sample preparation kit. The library was sequenced on an Illumina NextSeq 2K (P3 flow cell) using the 100-cycle kit (Read 1 = 50 bp, Read 2 = 80, Index 1□=□10). The data was aligned to a combined mouse and reovirus reference genome (described below) using the STAR Solo (version=2.7.9a) pipeline to derive a feature x bead barcode expression matrix.

### Slide-seq data preprocessing and analysis

Slide-seq count matrix and the position information for every bead barcode were loaded into an AnnData object using scanpy (v1.9.1). After filtering the beads with less than 50 transcripts detected and after removing genes detected in less than ten beads, we log-normalized the slide-seq expression data and computed principal components using highly variable genes (minimum dispersion = 0.2, minimum mean expression = 1.0). The transcriptomes were then clustered, and differential gene expression analysis (two-sided wilcoxon test) was performed to label bead clusters. Neighborhood enrichment permutation test was performed using Squidpy^50^ (v1.2.2). Cell2location^51^ (v0.1) was used for deconvolution of the Slide-seq transcriptomes using the scRNAseq as a reference. Genes in the reference were filtered with cell_count_cutoff=5, cell_percent_cutoff=0.03, and nonz_mean_cutoff=1.12 to select for highly expressed markers of rare cell types while removing most uninformative genes. Cell type signatures were determined using NB regression and used for spatial mapping of scRNAseq cell types on Slide-seq data with hyperparameters N_cells_per_location=1 and detection_alpha=20.

### Reference genome and annotation

*Mus musculus* genome and gene annotations (assembly: GRCm38) were downloaded from the Ensembl genome browser, and reovirus strain Type-1-Lang genome and gene annotations were downloaded and compiled from the NCBI browser. We have shared reovirus genome sequence and annotation files on figshare with the identifier https://doi.org/10.6084/m9.figshare.c.5726372. Genomes were processed using the Cell Ranger v-3.0.0 (10x Genomics) pipeline’s mkref command.

### Single-cell RNAseq data processing and visualization

Cells with fewer than 200 unique genes or more than 25 percent of transcripts aligning to mitochondrial genes were removed. After quality control, we captured 6596, 7096, and 3483 single-cell transcriptomes from mock-infected hearts, 5970, 5086, and 3453 single cell transcriptomes from reovirus wild-type (WT)-infected hearts, and 5354, 7462, and 3955 cells from reovirus mutant K287T-infected hearts at 4, 7, and 10 dpi respectively. The single-cell transcriptomes were log-transformed and normalized using the Scanpy package verison-1.8.1^52^. We used Scanpy to choose the highly variable genes with min_disp=0.5 and max_mean=3 thresholds. We then performed mean centering and scaling while regressing out total UMI counts, percent mitochondrial transcripts, S score, and G2M score, followed by principal component analysis (PCA) to reduce the dimensions of the data to the top 20 principal components (PCs). Uniform Manifold Approximation and Projection (UMAP) and the Nearest Neighbor (NN) graph were initialized in this PCA space using the first 20 PCs. The cells were then clustered using the Leiden method with multiple values of clustering resolution to get fine (resolution=0.5) and broad (resolution=0.3) celltype clusters. Cell-type-specific canonical gene markers along with differentially expressed genes (wilcoxon method) for each cluster were used to assign cell type labels. Normalized gene expression was visualized on DotPlots, UMAP plots, and Violin plots across cell type groups. A few cell type clusters representing cell states of the same cell type were grouped into broad cell type groups using cell type marker genes and then used for downstream analysis. Differential gene expression analysis (DGEA) was performed using the rank_gene_groups function in Scanpy with the Wilcoxon statistical method. All gene module scores were calculated using the score_genes function in scanpy.

### Reclustering and analysis of endothelial cells, T cells, fibroblasts, and cardiomyocytes

Normalized gene expression for a specific cell type group was extracted from the combined scRNA-seq dataset. We used Scanpy to reselect the highly variable genes within that cell type group with min_disp=0.5 and max_mean=3 thresholds. We then performed mean centering and scaling while regressing out total UMI counts, percent mitochondrial transcripts, S score, and G2M score, followed by principal component analysis (PCA) to reduce the dimensions of the data to the top 20 principal components (PCs). Uniform Manifold Approximation and Projection (UMAP) and the Nearest Neighbor (NN) graph were initialized in this PCA space using the first 20 PCs. The cells were then reclustered using the Leiden method (resolution=0.5 for endothelial cells, resolution=0.3 for T cells, resolution=0.2 for fibroblasts, and resolution=0.3 for cardiomyocytes) to get cell type subclusters. Differentially expressed genes (wilcoxon method) for each subcluster were then used to assign cell subtype labels. Subclusters representing doublets and expressing markers of multiple cell types were then removed from the analysis. Normalized gene expression for differentially expressed genes and genes of interest was visualized on DotPlots and UMAP plots across celltype subgroups. Differential gene expression analysis (DGEA) was performed using the rank_gene_groups function in Scanpy with the Wilcoxon statistical method. All gene module scores were calculated using the score_genes function in Scanpy.

### Spatial transcriptomics data processing, integration, analysis, and visualization

Spatial transcriptomics data from barcoded spatial spots from four heart sections were log-normalized using the Scanpy package (v1.8.1). Scanpy package was then used to select highly variable genes for spatial transcriptomics data with min_disp=0.5 and max_mean=3 thresholds. We then performed mean centering and scaling while regressing out total UMI counts, percent mitochondrial UMIs, S score, and G2M score, followed by PCA on the spot gene expression matrix, and reduced the dimensions of the data to the top 20 principal components. UMAP and the NN graph were initialized in this PCA space. The spots were then clustered using the Leiden method with multiple values of clustering resolution. The method returned spot clusters representing different tissue regions, which were then visualized on H&E images as spatial transcriptomics maps for individual samples to assign anatomical regions. Normalized gene expression was visualized on spatial transcriptomics maps for all tissue sections. Spot clusters representing the same tissue regions were grouped into broad anatomical region groups using marker genes and then used for downstream analysis. cell2location (version=0.1) deconvolution method compactible with scanpy and scvi-tools^53^ (v0.16.4) package was used for integration of spatial transcriptomics data with time-matched scRNA-seq data and cell type prediction values for spatial transcriptomics spots were estimated for the infected heart at 7 dpi. DGEA for anatomical regions was performed using the rank_gene_groups function in Scanpy with the Wilcoxon statistical method.

### Viral transcript sequencing data processing, filtering, and visualization

Enriched viral transcript data were aligned to a combined mouse and reovirus Type-1-Lang genome for all infected samples. Viral unique molecule (UMI) counts were taken from the combined expression matrices and added as metadata in the host gene expression data. Viral UMI counts in empty droplets, droplets with low-quality cells (< 200 host UMI counts), droplets with viable cells (>=200 host UMI counts) were sorted by viral UMI and visualized on a histogram to filter out the cell-free ambient viral RNA enriched in the hybridization protocol. Using the distribution of viral UMI counts in empty droplets, thresholds of two viral UMIs and five viral UMIs were used to identify infected cells in the heart and ileum respectively. Viral transcripts in the infected cells were then visualized on a DotPlot to determine viral tropism in tissues.

### Gene Ontology term enrichment analysis for scRNA-seq and spatial transcriptomics

Gene Ontology (GO) term enrichment analysis was performed on differentially expressed genes using gseapy (v0.10.4) wrapper package^54^. Differentially expressed genes (two-sided Wilcoxon test, log fold-change threshold = 2.0, p-value < 10^−4^ for scRNA-seq cells, and log fold-change threshold = 0.5, p-value < 10^−2^ for spatial transcriptomics spots) were selected and used for GO term enrichment analysis using GO_Biological_Processes_2021 gene sets in enrichr command^55^. The enriched GO terms of interest were selected and visualized on a BarPlot. The genes associated with GO terms of interested were used to calculate module scores using score_genes command in Scanpy.

### Sample preparation for RNA fluorescence in-situ hybridization (FISH), immunofluorescence, and histology

Whole hearts were isolated using aseptic technique and placed in ice cold sterile Hank’s Balanced Salt Solution and then blood was carefully removed by perfusing the hearts with fresh HBSS through the apex. Fresh tissues were immediately embedded in Optimal Cutting Compound (OCT) media and frozen in liquid nitrogen cooled isopentane, cut into 10 µm sections using a Thermo Scientific Microm 550 cryostat, and mounted on -20°C cooled histological glass slides which were then stored at -80°C until used.

### RNA fluorescence in-situ hybridization (FISH) split probe design and Signal Amplification using Hybridization Chain Reaction HCR-V3

Two-step hybridization strategy with split probe design and Hybridization Chain Reaction (HCR)-V3^56^ was used to label up to three transcripts in a single tissue section. Probes were designed using NCBI primer-blast which uses primer3 for designing internal hybridization oligo and BLASTn to check for binding specificity. We designed 20-21 bp primer pairs for an amplicon length of 40-42 bp (2 x primer length), primer melting temperature between 57°C and 63°C, and primer GC content between 35% and 65%. 7-10 sets of reverse complemented forward primers and reverse primers were then concatenated to flanking initiator sequence for HCR, ordered from Integrated DNA Technologies (IDT) with standard desalting purification (**Supp Data 4**). Split probes for each gene target, mixed and diluted in nuclease-free water to create a split probe pool stock solution at 10µM total probe concentration for every target. Hairpin pairs labeled with three different fluorophores namely Alexa-488, Alexa-546, and Alexa-647 (Molecular Instruments, **Supp Data 5**) were used for HCR V3.

### RNA fluorescence in-situ hybridization (FISH) experiments

Slides with tissue sections were then brought to room temperature until the OCT melts and were then immediately fixed in 4% paraformaldehyde for 12 minutes at room temperature. Post fixation, the sections were washed for 5 mins in 1x PBS twice, incubated for 1 hour in 70% ethanol for tissue permeabilization, washed again for 5 mins in 1x PBS, and then used for primary hybridization. Hybridization Buffer (HB) mix was prepared with 2x SSC, 5x of Denhart Solution, 10% Ethylene Carbonate, 10% Dextran Sulfate, 0.01% SDS, 1uM of probe pool mix per target for the hybridization reaction. 20 µl of HB mix (with probes) per section was then put on each slide to cover the tissue section, covered with parafilm, and incubated overnight at 37°C inside a humidifying chamber for primary hybridization. After primary hybridization, parafilm was removed and slides were washed in Hybridization Wash Buffer-1 (0.215M NaCl, 0.02M Tris HCl pH 7.5, and 0.005M EDTA) for 20-30 minutes at 48°C. Amplification Buffer (AB) mix was prepared with 2x SSC, 5x of Denhart Solution, 10% Dextran Sulfate, 0.01% SDS, 0.06µM of HCR hairpins for the amplification reaction. 2ul of each fluorophore labeled hairpins at 3µM corresponding to the target genes were mixed, incubated at 95°C for 1.5 minutes, covered in aluminum foil, and left to cool down at room temperature for 30 minutes to form hairpins before adding it to AB mix. 20 µl of AB mix per section was then put on each slide to cover the tissue section, covered with parafilm, and incubated overnight at room temperature in dark for signal amplification. After signal amplification, parafilm was removed, and slides were washed in 5x SSCT buffer twice for 30-40 minutes and then twice for 10 mins. The slides were then carefully cleaned with Kimwipe and treated with Ready Probes Auto-fluorescence Quenching Reagent Mix (Thermo Fisher, R37630) for 5 minutes and washed three times in 1X PBS. Last, tissue sections were then counter stained with DAPI for 10 minutes at room temperature, washed for 5 minutes in 1x PBS twice, excess PBS cleaned using Kimwipe, immediately mounted on coverslips using Slowfade antifade media, left overnight for treatment, and imaged the next day on a Zeiss Axio Observer Z1 Microscope using a Hamamatsu ORCA Fusion Gen III Scientific CMOS camera. smFISH images were shading corrected, stitched, rotated, thresholded, and exported as TIFF files using Zen 3.1 software (Blue edition).

### Immunofluorescence Assays

Slides with tissue sections were brought from -80°C freezer and heated for 1 minute at 37°C until the OCT melts and were then immediately dipped in prechilled methanol at -20°C for 30 minutes. After fixation, the slides were then rehydrated to Milli-Q water for 2 minutes and then washed twice in 1X PBS. Samples then underwent an antigen retrieval step via incubation in 1X citrate buffer for 10-15 minutes at 95°C. Samples were then permeabilized in 0.1% Triton X-100 in 1X PBS for fifteen minutes, washed three times in 0.05% Tween-20 in PBS (TBST), blocked for one hour at room temperature in blocking buffer (1% bovine serum albumin and 10% normal donkey serum in PBS. 20ul of primary antibodies diluted in antibody solution (1% bovine serum albumin in PBS) were then added on to the slides, covered with parafilm, then incubated in a humidifying chamber overnight at 4°C. Primary antibodies used were rabbit anti-reovirus VM1:VM6 polyclonal sera (1:30000), rat anti-Caspase1 monoclonal antibody (1:200, #14-9832-82, Invitrogen), rabbit anti-cleaved Caspase1 (Asp297) (1:200, #4199T, Cell Signaling Technology), and rabbit anti-cleaved Gasdermin D (Asp275) (1:200, #36425S, Cell Signaling Technology). Cleaved Caspase1 and cleaved Gasdermin D antibodies were purchased as a part of Pyroptosis Antibody Sampler Kit (#43811T, Cell Signaling Technology). After overnight primary incubation, samples were washed three times in PBS and then incubated in secondary antibodies diluted in blocking solution for two hours at room temperature. The secondary antibodies were donkey anti-rabbit alexa-488 (1:500, 711-545-152, Jackson Immuno Research), donkey anti-rabbit alexa-647 (1:500, 711-605-152, Jackson Immuno Research), and donkey anti-rat alexa-647 (1:500, ab150155, Abcam). Lastly, samples were washed thrice in PBS for 10 minutes with shaking, counter stained with DAPI, and mounted in Prolong antifade mounting media. Images were acquired on a Zeiss Axio Observer Z1 Microscope using a Hamamatsu ORCA Fusion Gen III Scientific CMOS camera. Immunostaining images were shading corrected, stitched, rotated, thresholded, and exported as TIFF files using Zen 3.1 software (Blue edition).

### Processing and quantification of Histology, RNA FISH, and immunofluorescence images

Image analysis and processing for histology, immunofluorescence, and RNA FISH images was done manually in Zen 3.1 software (Blue edition) and Fiji ImageJ. Whole heart Hematoxylin and Eosin (H&E) images were thresholded using non-linear adjustments (gamma = 0.45) applied across entire images using Zen 3.1 Blue software. For area quantifications from Hematoxylin and Eosin (H&E) stained histology images, 3-color RGB images were opened in ImageJ. The images were converted to greyscale 8-bit images and thresholded to detect the entire tissue section area. Sites of inflammation were manually selected for calculating the inflammation percentage in the tissue. For RNA FISH images, the images with DAPI counterstain channel were manually thresholded to segment nuclei. Holes in nuclei segmentation mask were files and morphological opening was performed to remove noise. The segmentation was enhanced using watershed algorithm followed by a morphological opening operation. For RNA FISH images, individual channels TIFF files exported from Zen 3.1 software were opened in ImageJ and converted to 8-bit images. Images were manually thresholded using linear adjustments (gamma = 1.0) applied across entire images to detect RNA-labelled cells and morphological opening was performed to remove noise. The nuclei and cells were counted in all images using the Analyze Particle function in ImageJ. For immunofluorescence images, individual channels were thresholded using linear adjustments (gamma = 1.0) applied across entire images. Thresholded images were loaded in Fiji ImageJ and converted to 8-bit images. The greyscale images for individual channels were then used to segment signal using same thresholds across all tissue sections to get selections for area quantifications. The tissue border was manually removed from the fluorescence channels when calculating the area of interest. Entire hearts were manually selected using DAPI channel to calculate total area of the tissue. Any changes to brightness and contrast were applied equally across the entire image for visibility of fluorescence signal.

## ACKNOWLEDGEMENTS

We would like to thank Peter Schweitzer and the Cornell Genomics Center for help with single-cell and spatial sequencing assays, Cornell Bioinformatics facility for assistance with bioinformatics, and Dr. Danica M. Sutherland from the lab of Dr. Terence S. Dermody at University of Pittsburgh for assistance with animal experiments and for providing the anti-reovirus polyclonal sera. We also thank the members of the Parker and De Vlaminck labs for many valuable discussions. This work was supported by R21AI144557 (to J.S.P. and I.D.V.), and DP2AI138242 (to I.D.V.). M.M. was supported by Distinguished Scholar Award from Center for Vertebrate Genomics, Cornell University. S.T.C. was supported by the National Institutes of Health and National Institute of Allergy and Infectious Diseases Award T32AI145821.

## AUTHOR CONTRIBUTIONS

M.M., J.S.P., and I.D.V. designed the study. M.M., M.M.H., and S.T.C. performed the animal experiments. M.M. and M.M.H. performed the scRNA-seq and spatial transcriptomics experiments. M.M., D.W.M., and M.F.Z.W. analyzed the data. M.M. performed histology, RNA FISH, and immunostaining experiments and analyzed the images. M.M., J.S.P., and I.D.V. wrote the manuscript. All authors provided feedback and comments.

## DATA AVAILABILITY

The authors declare that all sequencing data supporting the findings of this study have been deposited in NCBI’s Gene Expression Omnibus (GEO)^57^ with GEO series accession number <u>GSE189636</u>. Raw and processed H&E-stained tissue images and tissue-spot alignment files matched to spatial transcriptomics datasets have been made publicly available on figshare with identifier https://doi.org/10.6084/m9.figshare.c.5726372^58^. Scripts to reproduce the analysis presented in this study have been deposited on GitHub (https://github.com/madhavmantri/reovirus_induced_myocarditis).

## CONFLICTS

The authors declare no conflicts.

## Notes

### Competing Interest Statement

The authors have declared no competing interest.

### Summary of Updates

The updated manuscript file and supplementary imformation file contain smFISH and immunostaining results for an increased number of n's, new immunostaining experiments, new slide-seq data and analysis.

