## Supplement Material for "Spatiotemporal transcriptomics reveals pathogenesis of viral myocarditis"

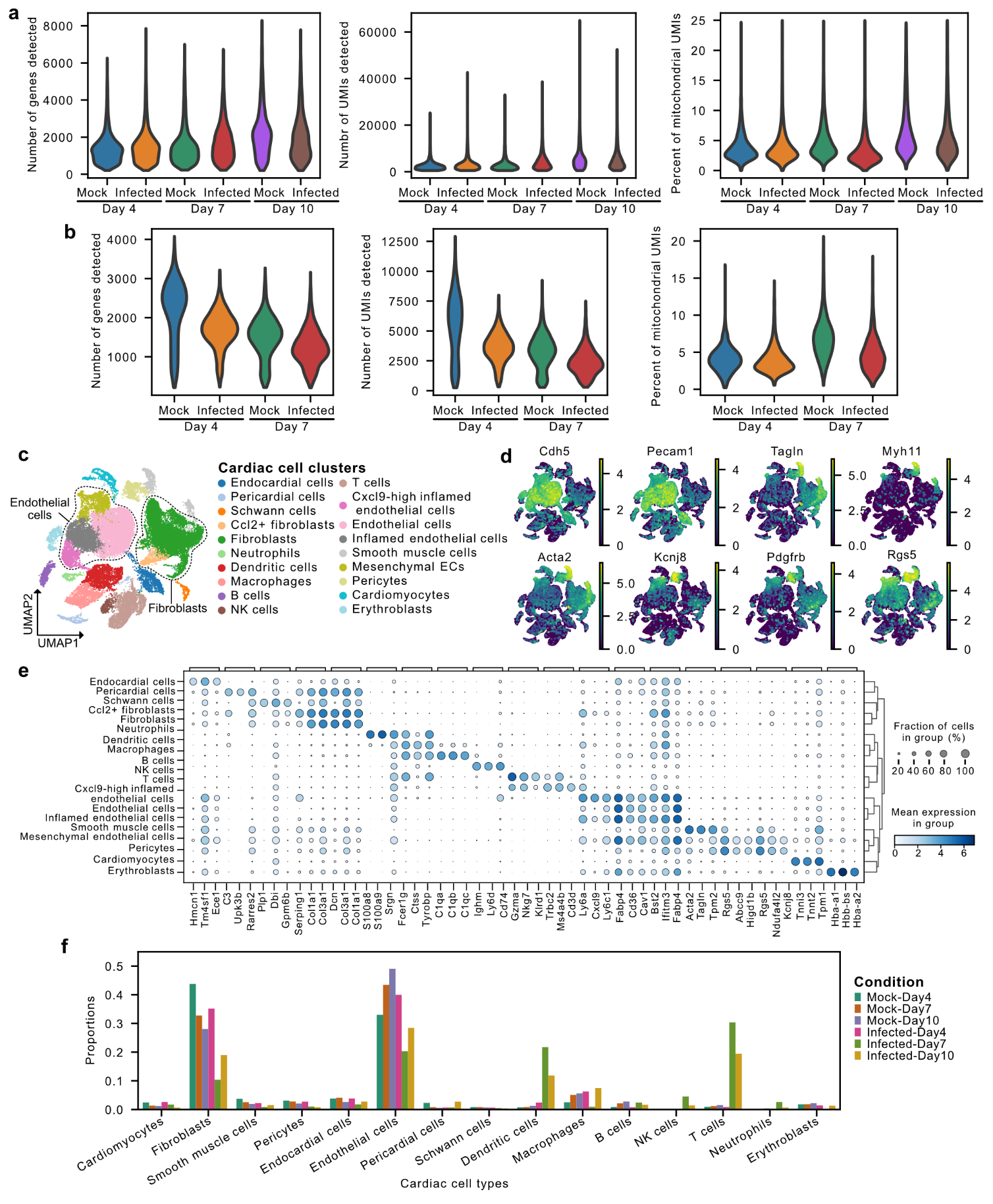

**Supplementary Figure 1: Single-cell and spatial transcriptomics of cardiac tissue from reovirus-infected neonatal mice. a)** Number of unique genes detected per cell (left), number of unique transcripts per cell (center), and percentage of mitochondrial transcripts (right) in cardiac scRNA-seq datasets from three stages after infection. **b)** Number of unique genes detected per cell (left), number of unique transcripts per cell (center), and percentage of transcripts from mitochondrial genes (right) in cardiac spatial transcriptomics datasets from two stages after infection. **c)** UMAP plot of 31,684 single-cell transcriptomes from mock-infected and reovirus-infected hearts at 4, 7, and 10 days post-infection (dpi), clustered by gene expression and colored by cardiac cell type. Dotted lines show the cardiac cell types being grouped as broad endothelial cells and fibroblast cells. **d)** scRNA-seq UMAP plots showing expression of endothelial markers *Cdh5* and *Pecam1*, smooth muscle cell-specific markers *Tagln*, *Myh11*, and *Acta2*, and pericyte markers *Pdgfrb*, *Kcnj8*, and *Rgs5* used to define the *Cdh5*+ *Kcnj8*+ *Pdgfrb*+ mesenchymal endothelial cells (Chen Qi et al. Nature Communications 2016). **e)** Top three differentially expressed genes (Wilcoxon test, log_2_ fold-change > 1.0 and p-value < 0.01) for cell types in heart scRNA-seq data. **f)** Bar plot showing the cell type composition changes in scRNA-seq datasets from reovirus-infected and mock-infected mice hearts.

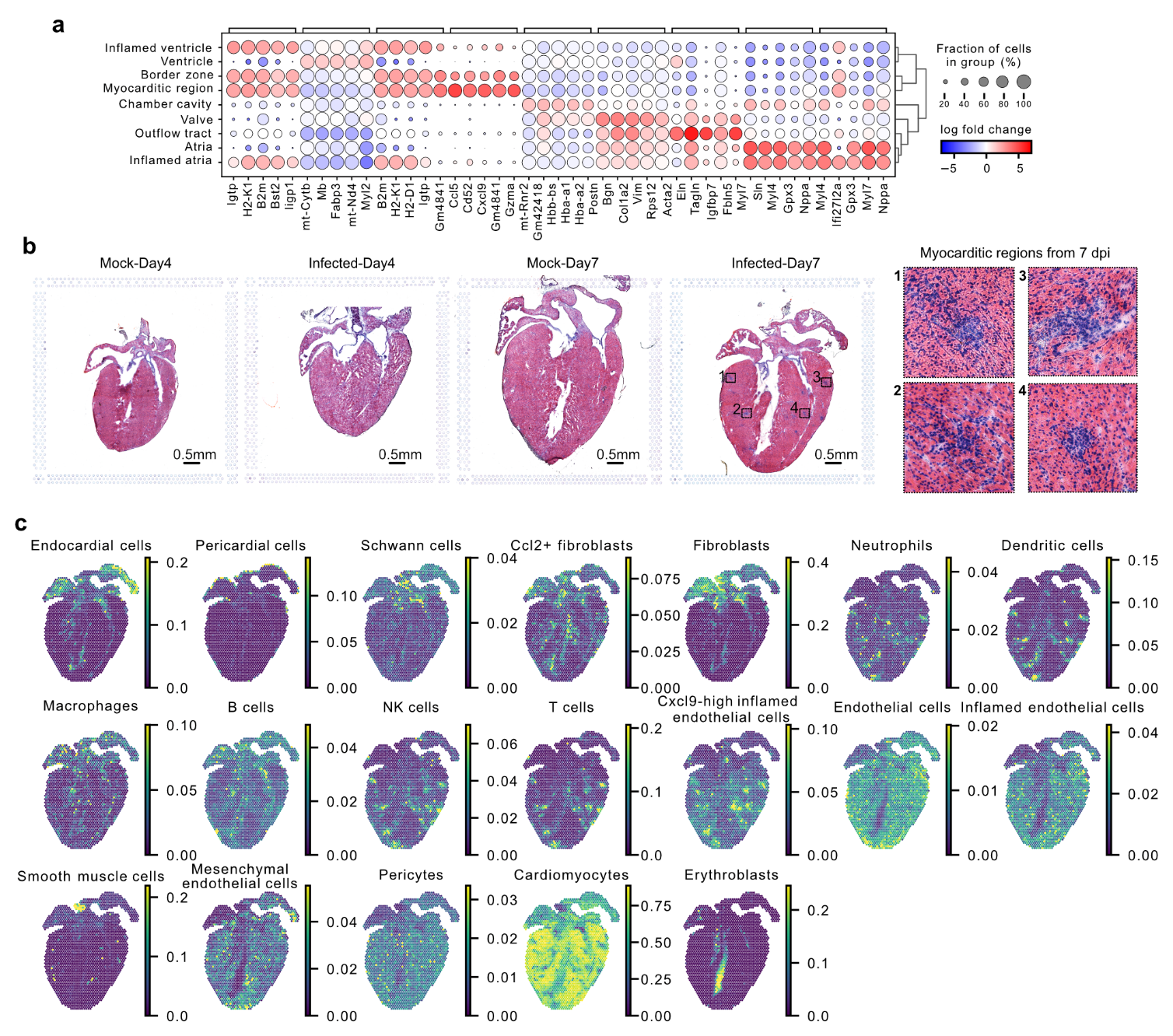

**Supplementary Figure 2: Single-cell and spatial transcriptomics of cardiac tissue from reovirus-infected neonatal mice. a)** Dot plot showing the top five differentially expressed genes (Wilcoxon test, log_2_ fold-change > 1.0 and p-value < 0.01) for spot clusters representing anatomical regions in in heart spatial transcriptomics datasets. **b)** Hematoxylin and Eosin (H&E) staining images on cardiac tissue sections used for spatial transcriptomics experiments. Insets show zoomed-in view of four myocarditic regions with inflammation in reovirus-infected heart at 7 days post infection (dpi). **c)** Spatial transcriptomics maps showing predicted cell type proportions for visium spatial transcriptomes from reovirus-infected heart at 7 dpi. scRNAseq data was used as a reference to perform cell type deconvolution using Cell2Location method.

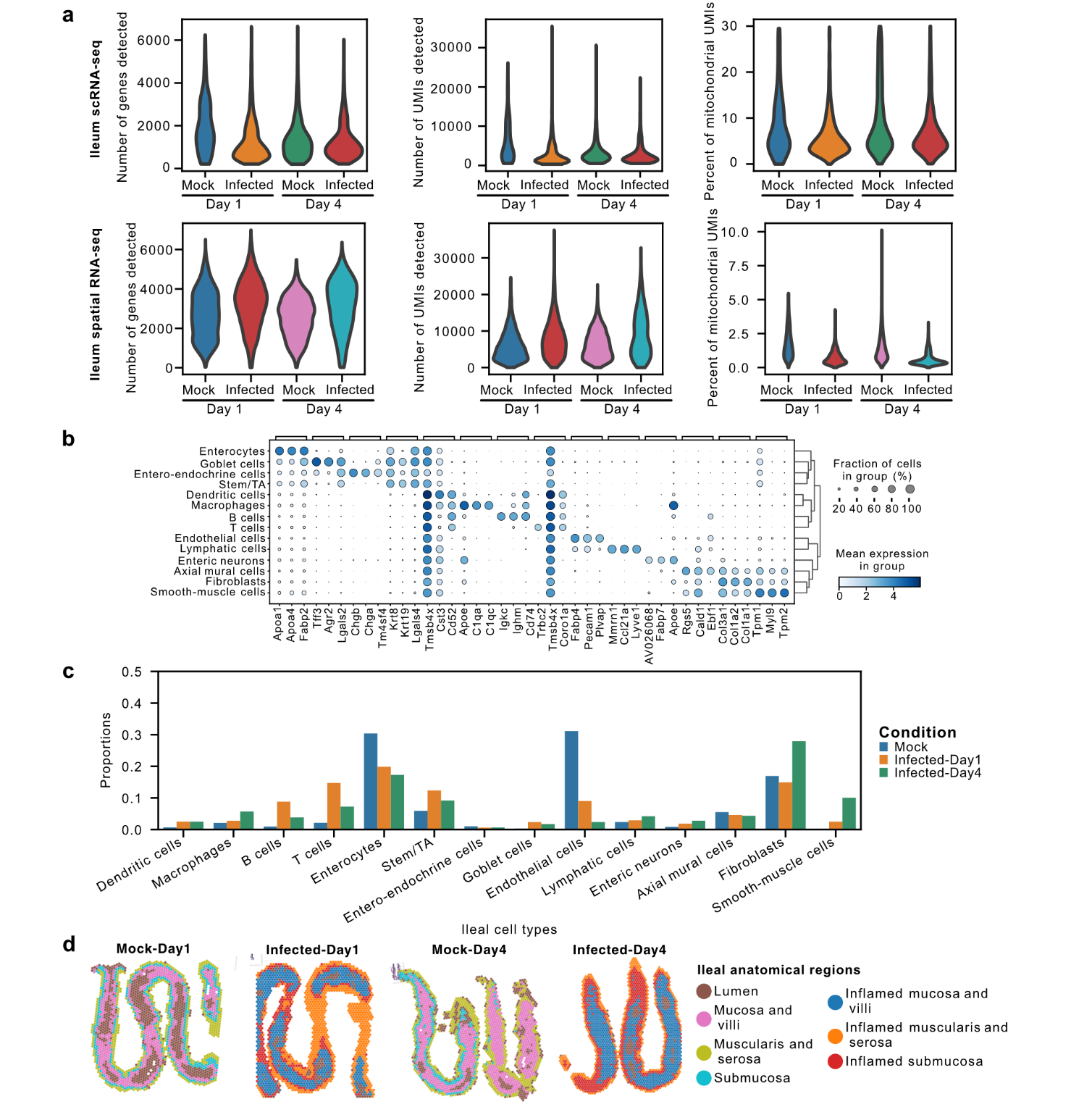

**Supplementary Figure 3: Single-cell and spatial transcriptomics of ileum tissue from reovirus-infected neonatal mice. a)** Number of unique genes detected per cell (left), number of unique transcripts per cell (center), and percentage of mitochondrial transcripts (right) in ileum scRNA-seq datasets (top row) and ileum spatial transcriptomics datasets (bottom row) from two stages after infection. **b)** Top-three differentially expressed genes (Wilcoxon test, log_2_ fold-change > 1.0 and p-value < 0.01) for cell types in ileum scRNA-seq data. **c)** Bar plot showing the cell type composition changes in scRNA-seq datasets from reovirus-infected and mock-infected mice in ileum tissue. Mock sample bar represents the mean cell type proportions for mock ileum samples at 1 and 4 dpi. **d)** Spatial transcriptomics map of ileum tissue sections from mock-infected and reovirus-infected mice at 1 and 4 dpi, colored by clusters representing tissue anatomical regions.

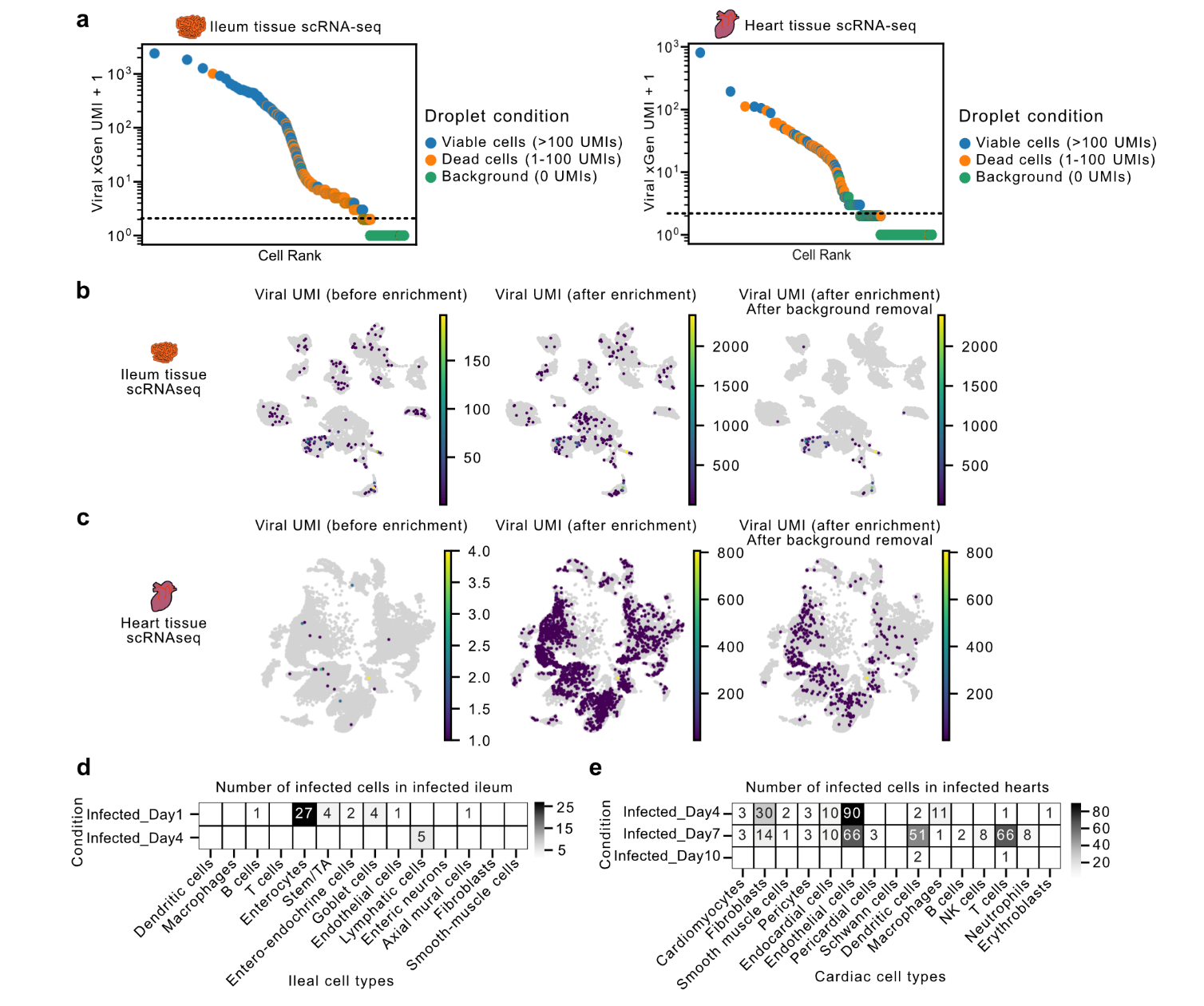

**Supplementary Figure 4: Enrichment of viral transcripts from single-cell transcriptomics libraries of ileum and heart tissue from reovirus-infected neonatal mice. a)** Knee plot showing viral UMI counts in the scRNA-seq droplets classified as either empty droplets or with viable/dead cells across ileum (left) and heart (right) scRNA-seq samples. The droplets were labelled using host gene UMI counts detected in scRNA-seq datasets. **b)** scRNA-seq UMAP plots showing total viral UMI counts per cell before xGen enrichment, after xGen viral transcript enrichment, and after removal of background signal on ileum samples. **c)** scRNA-seq UMAP plots showing total viral UMI counts per cell before xGen enrichment, after xGen viral transcript enrichment, and after removal of background signal on heart samples. **d)** Heatmaps showing counts of infected cells of different ileal cell types across two reovirus-infected ileum samples from 1 and 4 dpi. **e)** Heatmaps showing counts of infected cells of different cardiac cell types across three reovirus-infected heart samples from 4, 7, and 10 dpi.

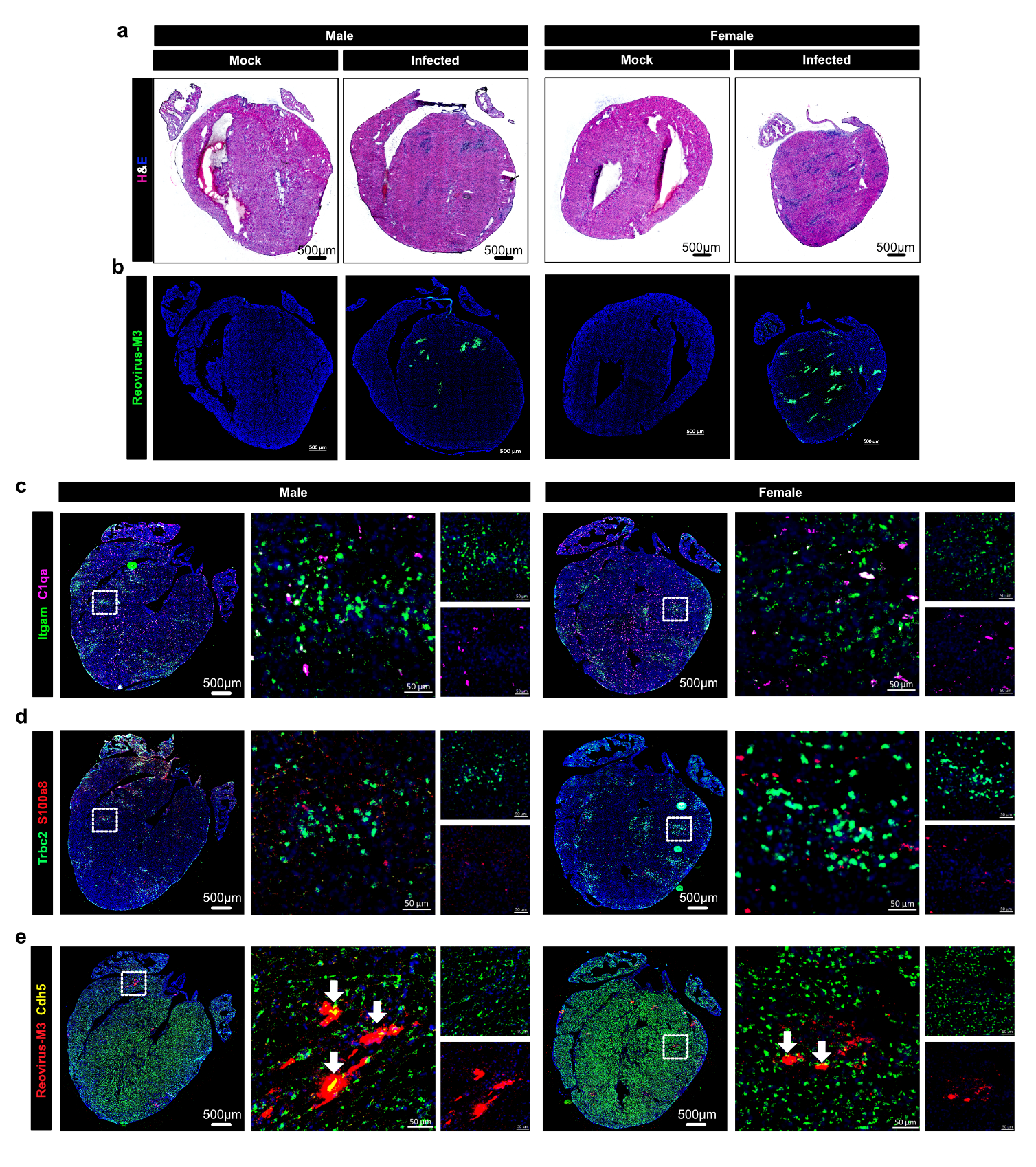

**Supplementary Figure 5: Imaging-based characterization of spatial distribution of different cardiac cell types in infected cardiac tissue. a)** Hematoxylin and Eosin (H&E) stained images of cardiac tissue sections from reovirus infected and mock infected mice at 7 dpi. **b)** Immunofluorescence staining of reovirus antigen in cardiac tissue sections from reovirus-infected and mock-infected hearts at 7 dpi. **c-d)** RNA FISH labeling of cell-type specific markers in reovirus-infected and mock-infected hearts at 7 dpi: macrophages (*Itgam*+ *C1qa*+), dendritic cells (*Itgam*+ *C1qa*-), neutrophils (*S100a8*), and T cells (*Trbc2*). **e)** RNA FISH labeling of endothelial cell marker Cdh5 and reovirus transcript (segment M3) in reovirus-infected and mock-infected hearts at 7 dpi. White arrows point at cell with reovirus transcripts. **b-e)** Representative images from 14 reovirus-infected (n=7 males and n=7 females) and six mock-infected (n=3 males and n=3 females) biological replicate mice.

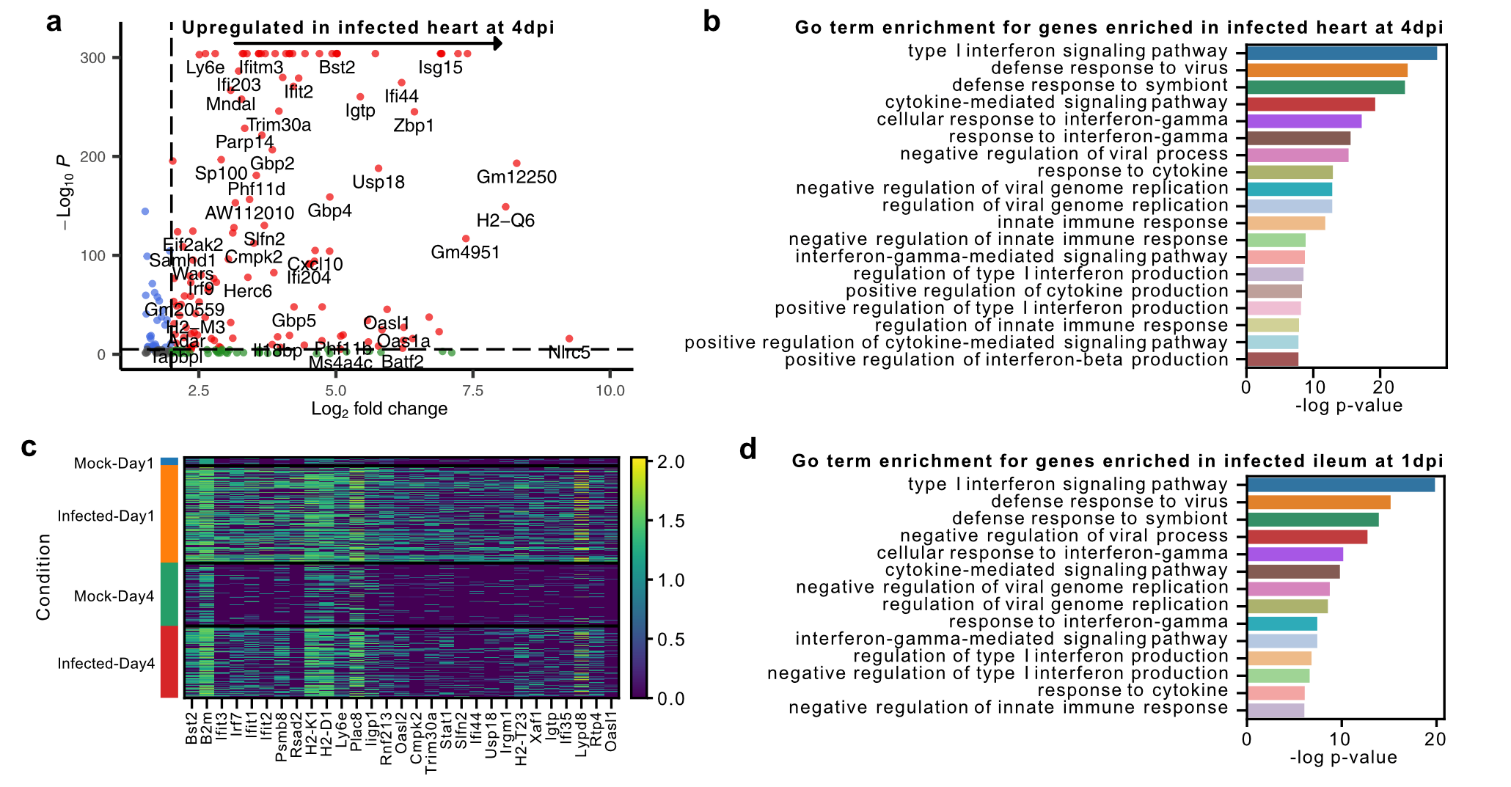

**Supplementary Figure 6: Innate immune response across cell types in heart and ileum tissue from reovirus-infected neonatal mice.** **a)** Volcano plot showing differentially expressed genes (two-sided Wilcoxon Rank-Sum test, -log2 fold change > 2.0 and p-value < 10-4) for reovirus-infected cardiac cells as compared to mock at 4 dpi. Dotted lines show the thresholds for significantly enriched genes (red). **b)** Top Gene Ontology (GO) terms for genes enriched in reovirus-infected cardiac cells as compared to mock at 4 dpi. **c)** Heatmap showing the expression of the 25 most upregulated genes in the reovirus-infected ileum as compared to mock at 1 dpi. **d)** Top GO terms for genes enriched in reovirus infected cells ileum as compared to mock-infected ileum at 1 dpi.

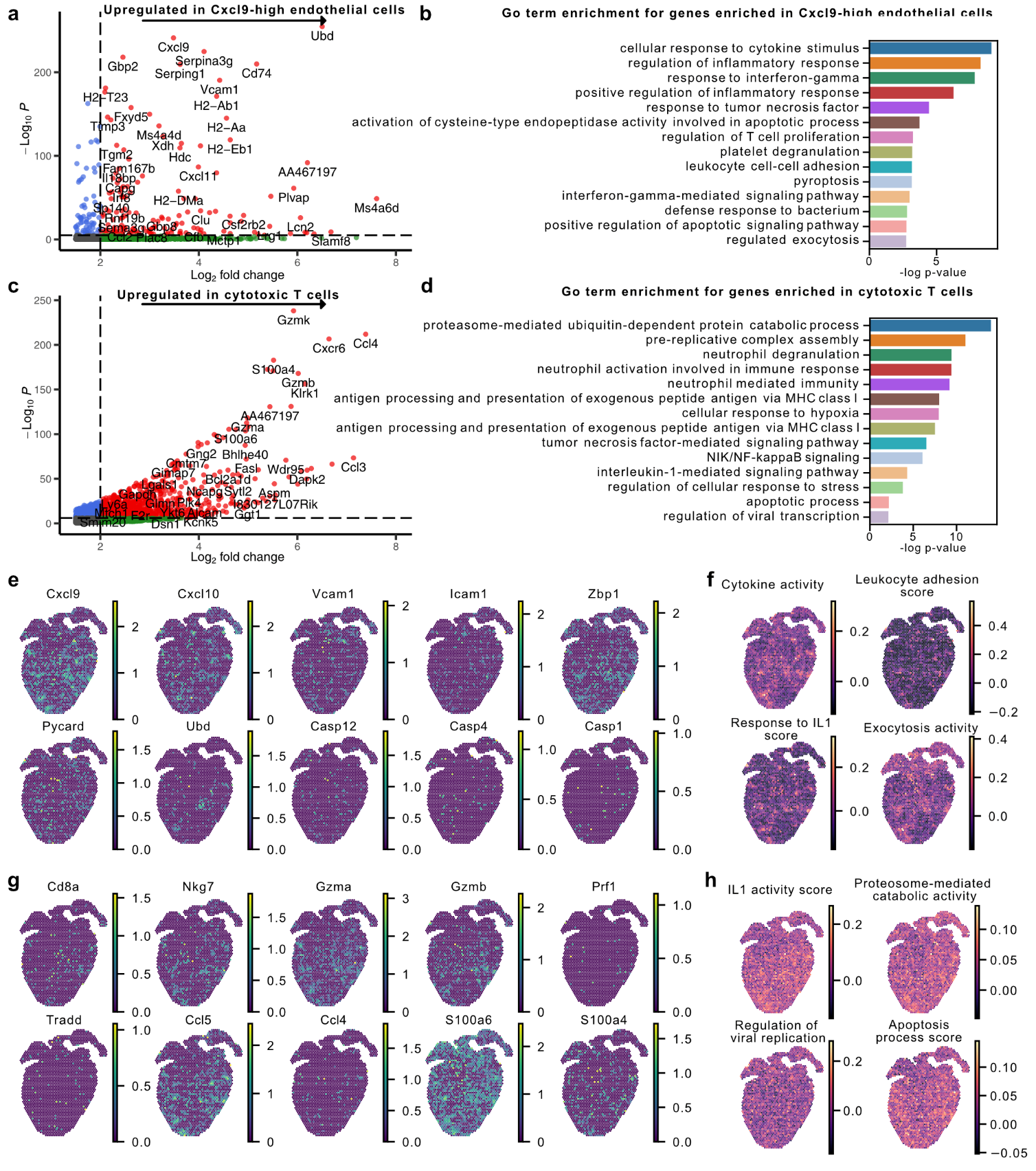

**Supplementary Figure 7: Transcriptional gene signatures and programs for Cxcl9-high endothelial cells and cytotoxic T cells found within the myocarditic tissue. a)** Volcano plot showing differentially expressed genes (two-sided Wilcoxon Rank-Sum test, -log2 fold change > 2.0 and p-value < 10-4) upregulated in *Cxcl9*-high inflamed endothelial cells from heart at 7 dpi. Dotted lines show the thresholds for significantly enriched genes (red). **b)** Top Gene Ontology (GO) terms of interest enriched for genes upregulated in *Cxcl9*-high endothelial cells in myocarditic heart at 7 dpi. **c)** Volcano plot showing differentially expressed genes (two-sided Wilcoxon Rank-Sum test, -log2 fold change > 2.0 and p-value < 10-4) upregulated in cytotoxic T cells at 7 and 10 dpi. Dotted lines show thresholds for significantly enriched genes (red). **d)** Top GO terms of interest enriched for genes upregulated in cytotoxic T cells in myocarditic hearts at 7 and 10 dpi. **e-h)** Spatial transcriptomics maps of cardiac tissue sections from reovirus-infected mice pups at 7 dpi showing: **e)** The expression of genes enriched in *Cxcl9*-high inflamed endothelial cells. **f)** Gene module scores for four GO terms of interest enriched in *Cxcl9*-high inflamed endothelial cells. **g)** The expression of genes enriched in cytotoxic T cells from myocarditic heart. **h)** Gene module scores calculated for four GO terms of interest enriched in cytotoxic T cells from myocarditic heart.

s
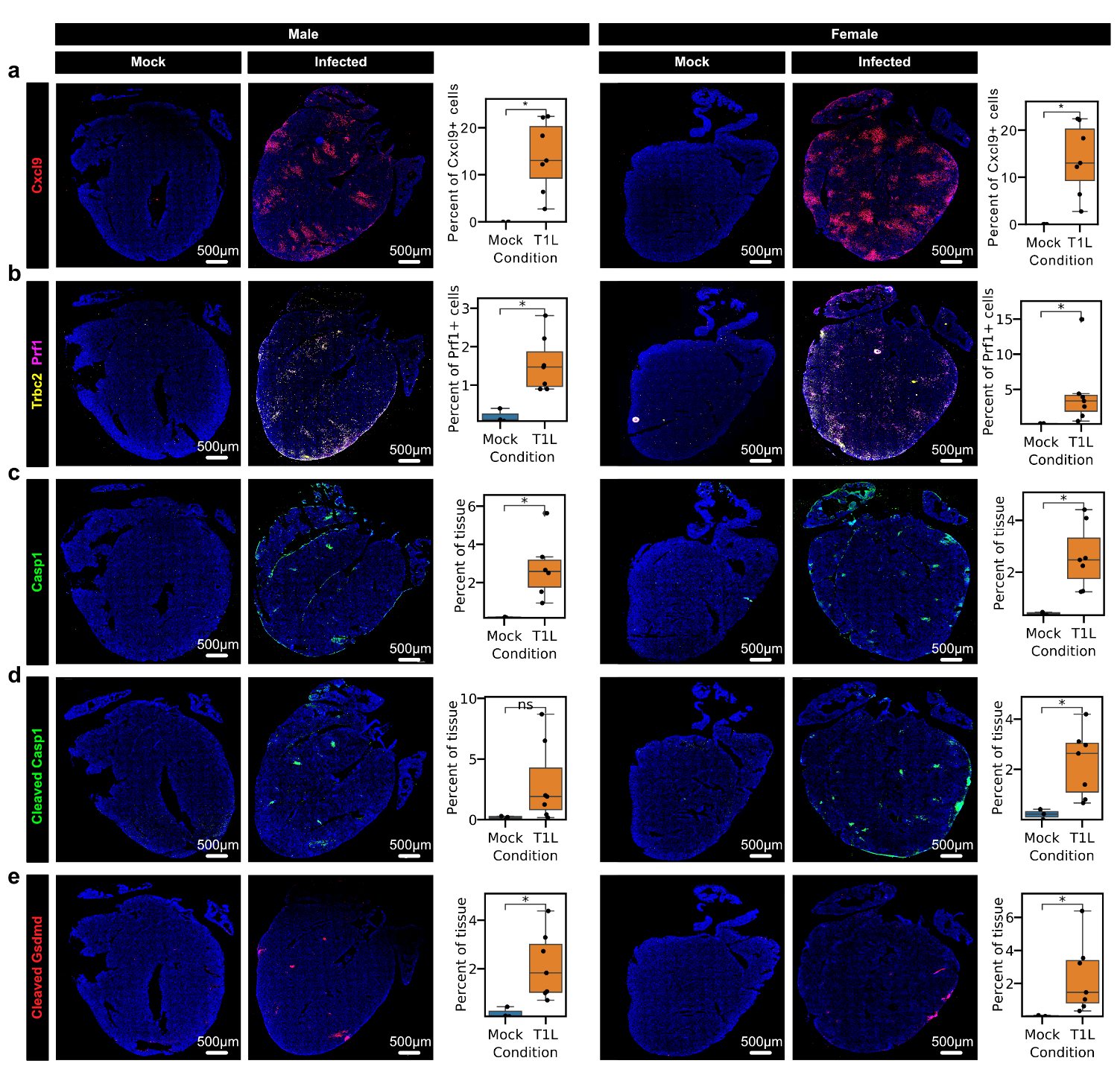

**Supplementary Figure 8: Imaging-based characterization of *Cxcl9*-high endothelial cells, cytotoxic T cells, and pyroptosis in myocarditic tissue. a)** RNA FISH labelling of *Cxcl9* transcripts in *Cdh5*+ endothelial cells in reovirus-infected and mock infected hearts at 7 dpi. **b)** RNA FISH labelling of *Prf1* transcripts in *Trbc2*+ T cells in reovirus-infected and mock-infected hearts, confirming the recruitment of cytotoxic T cells in myocarditic tissue (bottom row). **c-e)** Immunostaining of protein markers for pyroptosis activity: **c)** Casp1 protein (Pro-caspase1 and cleaved Caspase1) **d)** cleaved Caspase1 protein (only Casp1 p20 subunit) and **e)** Cleaved Gasdermin D (Gsdmd N terminus fragment) in reovirus-infected and mock-infected hearts at 7 dpi. **b-e)** Representative images from 14 reovirus-infected mice (n=7 males and n=7 females) and six mock-infected mice (n=3 males and n=3 females). Immunofluorescence signal from reovirus-infected hearts was compared to mock-infected hearts using two-sided Wilcoxon statistical test. p-value annotation legend: ns: p <= 1.00e+00, *: 1.00e-02 < p <= 5.00e-02, **: 1.00e-03 < p <= 1.00e-02, ***: 1.00e-04 < p <= 1.00e-03, ****: p <= 1.00e-04.

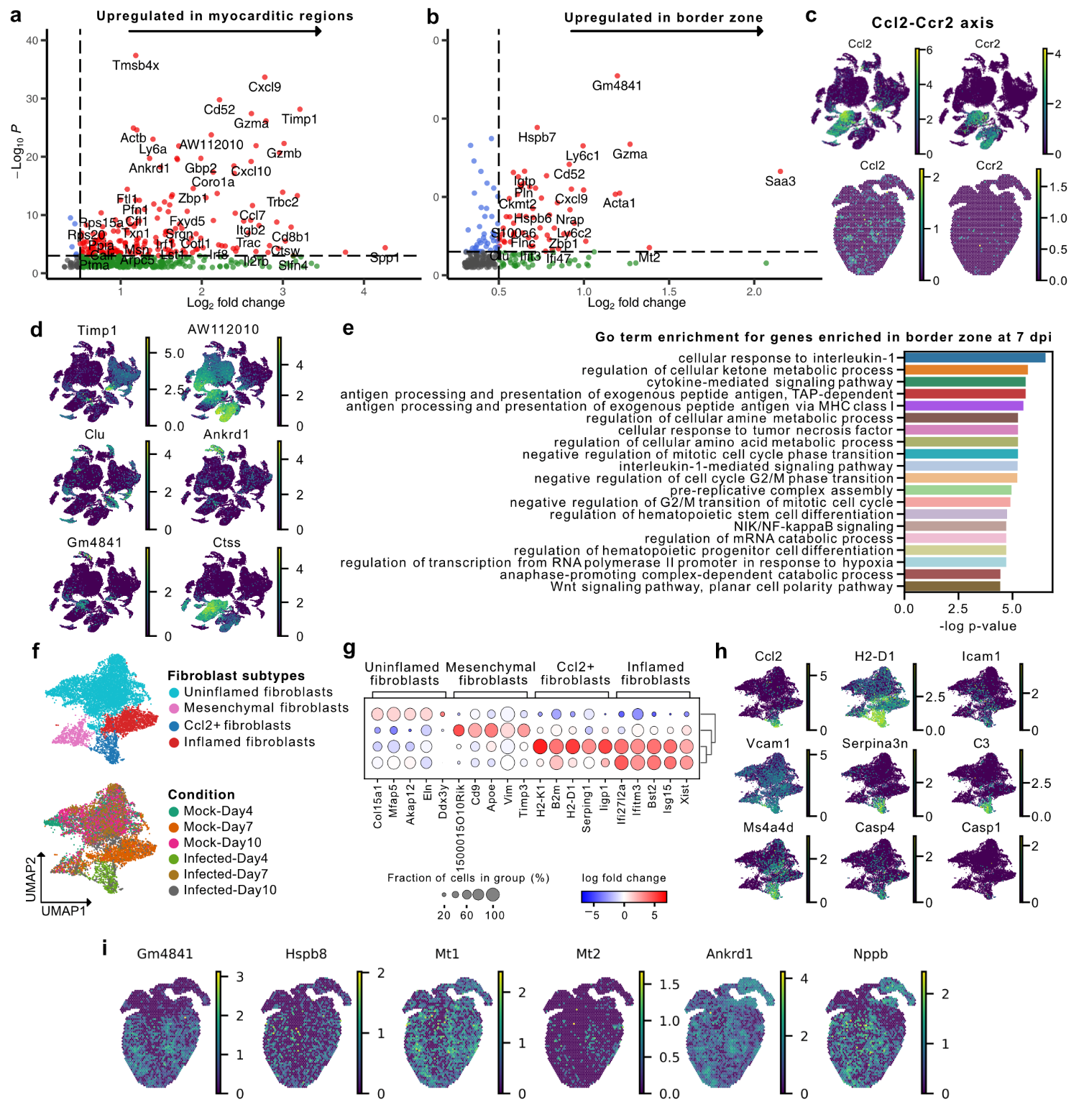

**Supplementary Figure 9: Cell-type-specific gene expression in myocarditic tissue and characterization of inflamed fibroblast phenotype found within the myocarditic tissue. a)** Volcano plot showing differentially expressed genes (two-sided Wilcoxon Rank-Sum test, -log_2_ fold change > 0.5 and p-value < 10^-2^) upregulated in myocarditic regions in heart at 7 dpi as defined by unsupervised clustering on spatial transcriptomes. Dotted lines show the thresholds for significantly enriched genes (red). **b)** Volcano plot showing differentially expressed genes (two-sided Wilcoxon Rank-Sum test, -log_2_ fold change > 0.5 and p-value < 10^-2^) upregulated in the border zone of the infected heart at 7 dpi, as defined by unsupervised clustering on spatial transcriptomes. Dotted lines represent thresholds for significantly enriched genes (red). **c)** UMAP plot for heart scRNA-seq cells and spatial transcriptomic maps for reovirus-infected heart at 7 dpi showing the expression of *Ccl2* ligand and *Ccr2* receptor. **d)** scRNA-seq UMAP plots showing the expression of six genes of interest enriched in myocarditic regions and the border zone. **e)** Top GO terms of interest for genes upregulated in *Cxcl9*-high endothelial cells in myocarditic heart at 7 dpi. **f)** UMAP plot of 9,192 fibroblast cell transcriptomes from mock-infected and reovirus-infected hearts at 4, 7, and 10 dpi colored by fibroblast cell subtypes (phenotypes) (top) and condition (bottom). **g)** Heatmap showing top-five differentially expressed genes (Wilcoxon test, log_2_ fold-change > 1.0 and p-value < 0.01) for fibroblast cell subtypes. **h)** UMAP plot showing the expression of genes upregulated in *Ccl2*+ fibroblast cells. **i)** Spatial transcriptomic maps for myocarditic heart at 7 dpi showing the expression of six myocyte-specific genes upregulated in the border zone.

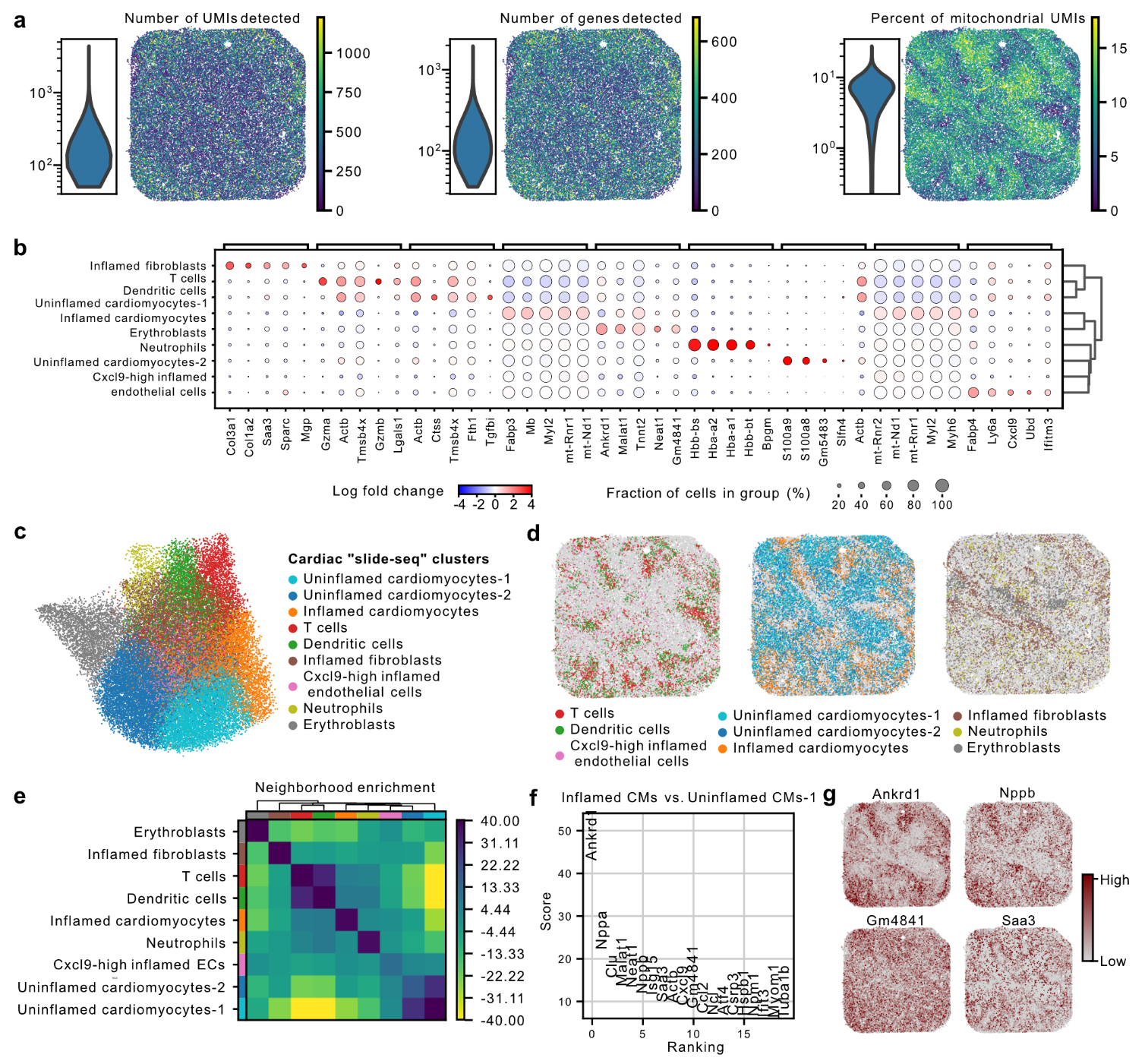

**Supplementary Figure 10: High-resolution slide-seq spatial transcriptomics of cardiac tissue from reovirus-infected neonatal mice. a)** Number of unique UMIs detected per cell (left), number of unique genes detected per cell (center), and percentage of mitochondrial transcripts (right) in slide-seq datasets from reovirus infected heart at 7 dpi. **b)** Heatmap showing top-five differentially expressed genes (Wilcoxon test, log_2_ fold-change > 1.0 and p-value < 0.01) for slide-seq spatial transcriptomics clusters. **c)** UMAP plot of > 40,000 slide-seq spatial transcriptomes from reovirus-infected heart at 7 dpi, clustered by gene expression and colored by putative cardiac cell types based on differential gene expression and marker analysis. **d)** Slide-seq spatial transcriptomics maps showing three slide-seq clusters at a time. **e)** Heatmap of permutation test scores for neighborhood enrichment of slide-seq clusters. Enrichment scores reflect enrichment of spatial proximity of slide-seq clusters. **f)** Rank plot showing differential gene expression results between inflamed and uninflamed cardiomyocytes in slide-seq spatial transcriptomics data (two-sided Wilcoxon test, log-foldchange > 1.0 and p-value < 0.01). **g)** Slide-seq spatial transcriptomics maps showing the expression of inflamed cardiomyocyte specific genes.

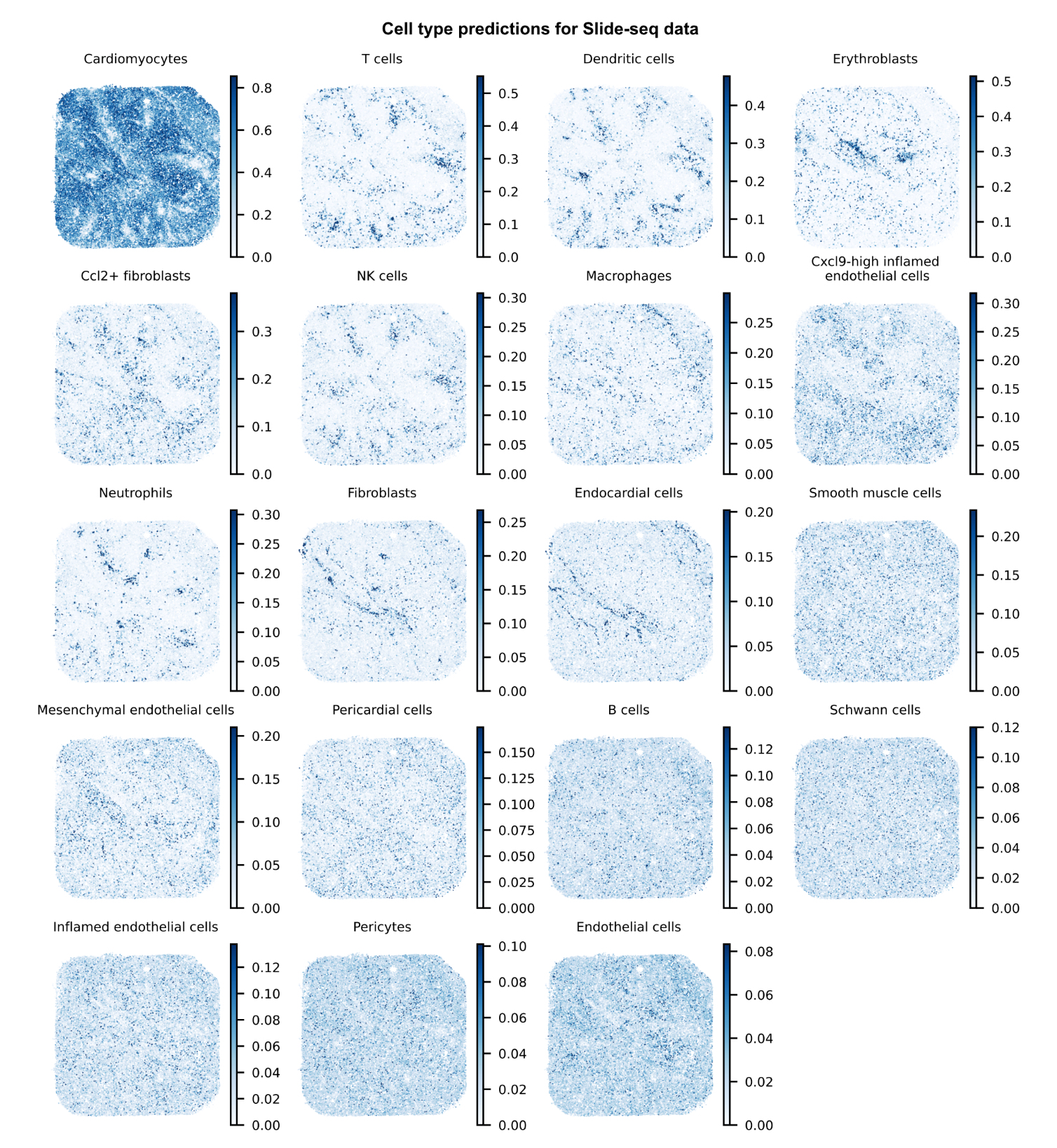

**Supplementary Figure 11:** Spatial transcriptomics maps of predicted cell type proportions for slide-seq spatial transcriptomes measured for a reovirus-infected heart at 7 dpi. The scRNAseq data in Figure of the main text was used as a reference to perform cell type deconvolution (cell2location method).

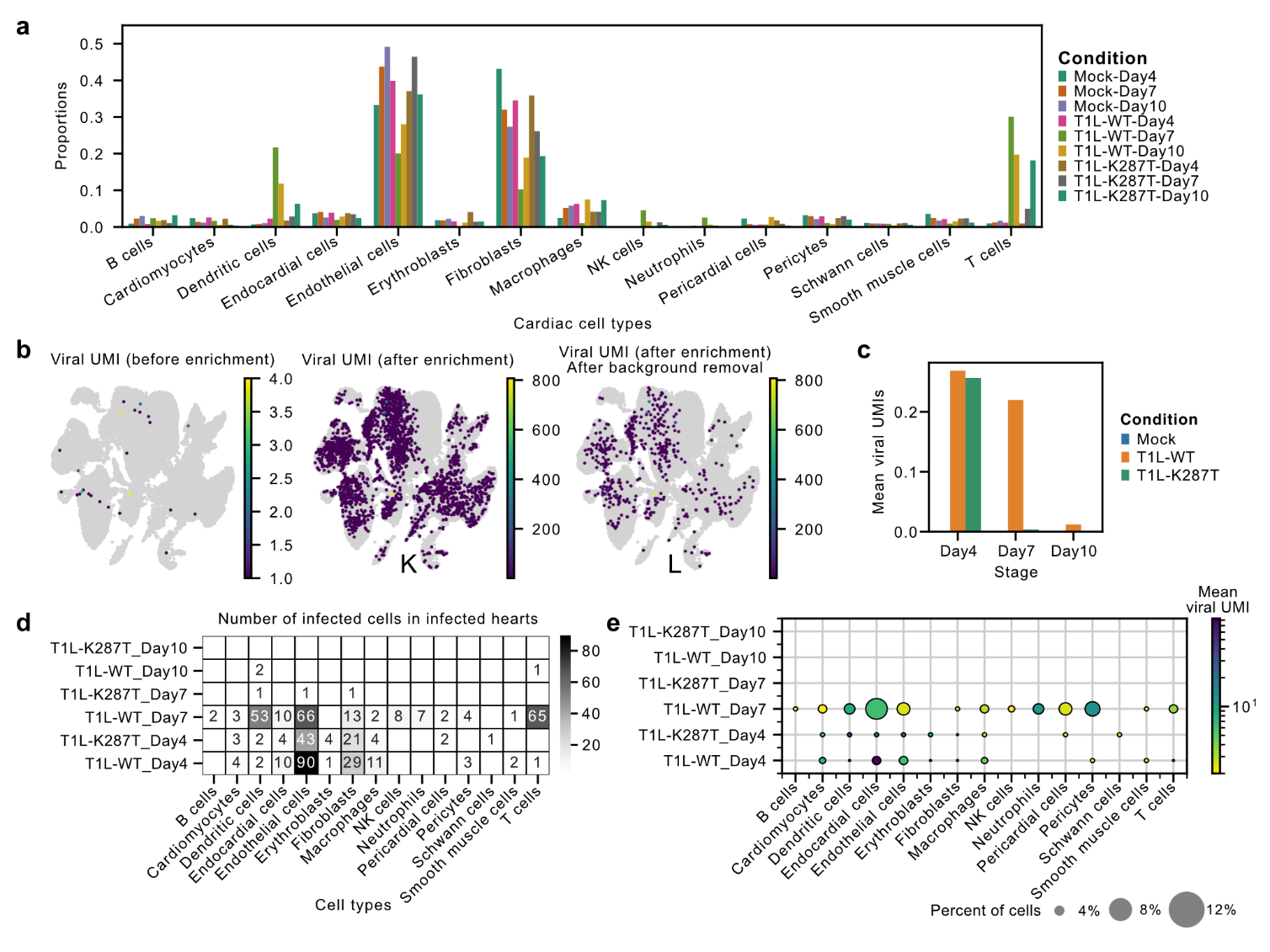
 **Supplementary Figure 12: Differences in cell type composition and viral tropism between scRNA-seq cells from reovirus WT and K287T mutant-infected neonatal mice. a)** Bar plot showing the cell type composition changes in scRNA-seq datasets from reovirus WT and K287T mutant-infected cardiac tissue. **b)** scRNA-seq UMAP plots showing total viral UMI counts per cell before xGen enrichment, after xGen viral transcript enrichment, and after removal of background signal on heart samples. **c)** Bar plot showing mean viral transcript count (UMIs) across stages for reovirus WT- and K287T mutant- infected hearts. **d)** Heatmaps showing counts of infected cells of different cardiac cell types across reovirus WT and K287T mutant-infected heart samples. **e)** Dot plot showing the percentage of cells with non-zero viral transcripts and the mean viral transcript counts (UMIs) across cell types.
